# A transcriptional regulatory circuit for the photosynthetic acclimation of microalgae to carbon dioxide limitation

**DOI:** 10.1101/2020.07.09.195545

**Authors:** Olga Blifernez-Klassen, Hanna Berger, Birgit Gerlinde Katharina Mittmann, Viktor Klassen, Louise Schelletter, Tatjana Buchholz, Thomas Baier, Maryna Soleimani, Lutz Wobbe, Olaf Kruse

**Affiliations:** Algae Biotechnology and Bioenergy, Bielefeld University, Faculty of Biology, Center for Biotechnology (CeBiTec), Universitätsstrasse 27, 33615, Bielefeld, Germany

**Author notes:** Die Blattmacher GmbH, Friedrichstraße 153a, 10117 Berlin (Hanna Berger). **To whom correspondence should be addressed:** Olaf Kruse, Bielefeld University, Faculty of Biology, Center for Biotechnology (CeBiTec), Universitätsstrasse 27, 33615 Bielefeld, Germany. The author responsible for distribution of materials integral to the findings presented in this article in accordance with the policy described in the Instructions for Authors (www.plantcell.org) is: Olga Blifernez-Klassen.

**Keywords:** Light-harvesting antenna, transcription factor, carbon dioxide responsive cis-regulatory elements, NAB1, *Chlamydomonas reinhardtii*

## Abstract

In green microalgae, prolonged exposure to inorganic carbon depletion requires long-term acclimation responses, based on a modulated expression of genes and adjusting photosynthetic activity to the prevailing supply of carbon dioxide. Here, we depict a microalgal regulatory cycle, adjusting the light-harvesting capacity at PSII to the prevailing supply of carbon dioxide in *Chlamydomonas reinhardtii*. It engages a newly identified low carbon dioxide response factor (LCRF), which belongs to the Squamosa promoter binding protein (SBP) family of transcription factors, and the previously characterized cytosolic translation repressor NAB1. LCRF combines a DNA-binding SBP domain with a conserved domain for protein-protein interactions and transcription of the *LCRF* gene is rapidly induced by carbon dioxide depletion. LCRF activates transcription of the *NAB1* gene by specifically binding to tetranucleotide motifs present in its promoter. Accumulation of the NAB1 protein enhances translational repression of its prime target mRNA, encoding the PSII-associated major light-harvesting protein LHCBM6. The resulting reduction of the PSII antenna size helps maintaining a low excitation during the prevailing carbon dioxide limitation. Analyses of low carbon dioxide acclimation in nuclear insertion mutants devoid of a functional *LCRF* gene confirm the essentiality of this novel transcription factor for the regulatory circuit.

## INTRODUCTION

In photosynthetic organisms, control of light-harvesting is a key component of acclimation mechanisms that adjust photon capture capacity to the prevailing external condition. A sudden drop in CO_2_ availability causes an increased excitation pressure at PSII, which is deleterious, if not rapidly relieved by short-term acclimation mechanisms. This increase in excitation pressure results from an over-reduction of the photosynthetic electron transport chain (for review see (Wobbe et al., 2016)), which is in turn caused by the reduced consumption of NADPH and ATP, the products of photosynthetic light reactions, by the Calvin-Benson-Bassham cycle. As an immediate response to the onset of high excitation pressure, non-photochemical quenching (NPQ) mechanisms are activated (Allorent et al., 2013). Among them are state transitions, which represent the predominant fast mechanism that reduces PSII excitation pressure in response to carbon dioxide depletion in *Chlamydomonas reinhardtii* (Bulté et al., 1990; Iwai et al., 2007; Lucker and Kramer, 2013; Takahashi et al., 2013) and are triggered by an over-reduced plastoquinone pool, activating LHCII phosphorylation and a diminished light absorption at PSII (Goldschmidt-Clermont and Bassi, 2015).

However, during prolonged carbon dioxide limitation, the LHC state II transition is reversed (Iwai et al., 2007), indicating that excitation pressure relief based on state transitions is replaced by long-term responses, like functional antenna size reduction (Spalding et al., 1984).

In a previous study, we could show that this reduction in functional antenna size occurs via LHCBM translation repression and that this can efficiently relieve excitation pressure at PSII under prolonged carbon dioxide depletion. The increased accumulation of the translation repressor NAB1 emerged as a key component within this response, and the application of a photosynthetic electron transfer (PET) inhibitor indicated that signals emerging from the chloroplast control nuclear *NAB1* promoter activity. In consequence, regulatory elements are presumably encoded in the promoter sequence, allowing an activation of *NAB1* transcription under carbon dioxide limitation (Berger et al., 2014).

In agreement with previous work (Berger et al., 2014), which clearly demonstrated an activation of the *NAB1* promoter by carbon dioxide depletion via reporter assays, a transcriptome study revealed changes in *NAB1* transcript abundance dependent on carbon dioxide supply (Winck et al., 2013).

Although nuclear gene expression modulation in response to carbon dioxide depletion is evident (Fang et al., 2012; Winck et al., 2013), information on the molecular mechanisms underlying these transcriptomic changes is limited (Wang et al., 2015). In general, only a small number of nuclear *Chlamydomona*s promoters have already been studied in the context of photosynthetic acclimation responses (Sawyer et al., 2015; Maruyama et al., 2014; Winck et al., 2013; Shao et al., 2008).

In this study, we systematically analyzed the *NAB1* upstream region regarding the location of *cis*-regulatory elements and exploited an identified minimal promoter, conferring carbon dioxide-responsiveness, within yeast-one-hybrid analyses, which enabled the identification of a novel transcription factor LCRF. Further, we show that LCRF is essential for the induction of NAB1 gene expression modulation in response to carbon dioxide limitation and hence antenna size control.

## RESULTS

### The transcription factor LCRF specifically binds to motifs present in the promoter of the *NAB1* gene

Our previous study demonstrated that the activity of the *NAB1* promoter is induced by carbon dioxide deprivation (Berger et al., 2014). To characterize the *NAB1* upstream region in more detail and to identify the *cis*-regulatory elements mediating an induction following carbon dioxide deprivation, the precise transcription start site was first determined by a modified “new 5’RACE” experiment (Scotto-Lavino et al., 2006). Cloning and sequencing of a 5’ full length cDNA demonstrated that transcription starts 102 bp upstream of the start codon of the *NAB1* coding sequence (Phytozome v12.1, Cre06.g268600). Under the conditions examined, no fragment larger than 102 bp could be detected, strongly suggesting that this marks the end of the 5’UTR of the *NAB1* gene. To confirm this result, the approximate length of the 5’UTR was mapped by PCR, taking complementary (cDNA) or genomic DNA (gDNA) as templates (Figure 1A). A reverse primer binding close to the translation start site was combined with three distinct forward primers (Supplementary Table S1; mapping 5’UTR), designed to bind 100, 147 and 265 bp upstream of the translation start (Figure 1A, ATG).

**Figure 1:**
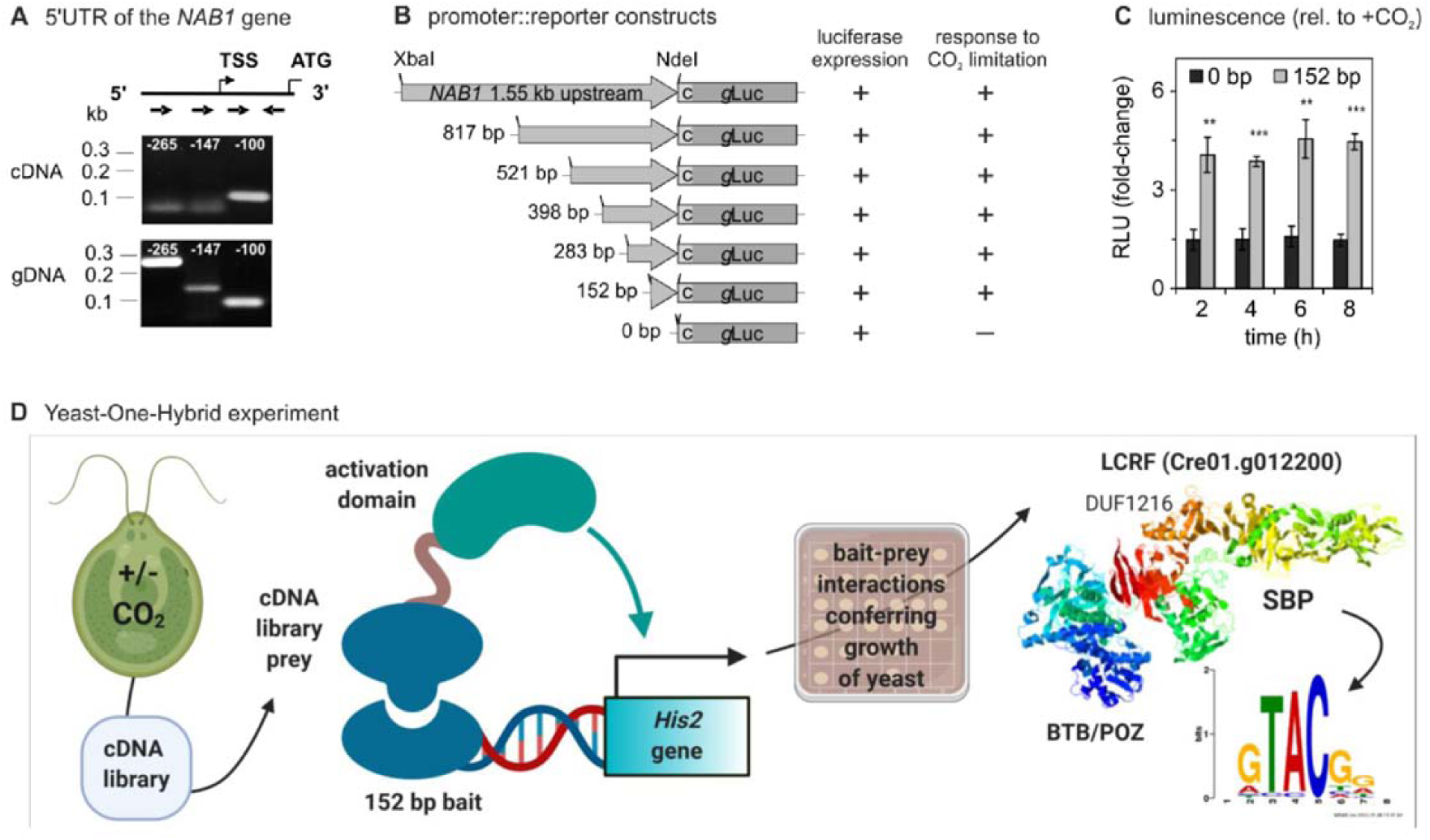
A 152 bp sequence is sufficient to confer carbon dioxide-responsiveness to the *NAB1* promoter. **(A)** Mapping of the 5’UTR of the *NAB1* gene. PCR was conducted with either genomic DNA (gDNA) or complementary DNA (cDNA), resulting from reverse transcription, as the templates. Distinct *NAB1* promoter-specific primers (forward primers binding 100, 147 and 265 bp upstream of the translation start; upper panel) were used and PCR products separated in 2% agarose gels prior to staining with SYBR Safe. **(B)** Promoter-bashing analysis performed with a sequence beginning 1.55 kb upstream and extending to the *NAB1* start codon. The full length and several truncated constructs were analyzed regarding their ability to drive expression of a *Gaussia* luciferase (gLuc) reporter and their responsiveness to carbon dioxide limitation (from 3% (v/v) CO_2_ to air levels) (see also Figure S1). **(C)** Luminescence assay to analyze expression induction following carbon dioxide deprivation in transformants containing a stably integrated *gLuc* reporter either driven by a 152 bp minimal promoter sequence or being devoid of a promoter (0 bp control). For each construct, the luminescence determined under CO_2_-replete conditions (3% (v/v) CO_2_) at t_0_ was set to 1. Error bars represent the standard error derived from experiments using three distinct cell lines per construct and include the mean values from three biological with three technical replicates per cell line (SEM, n=3). Asterisks represent p-values as determined via Student’s t-test (* ≤ 0.05, ** ≤ 0.01, *** ≤ 0.001). **(D)** Depiction of the workflow of the Yeast-One-Hybrid experiment conducted to identify candidate proteins binding the 152 bp fragment of the *NAB1* promoter. The shown LCRF protein model was predicted using iTASSER server (C-score 0.59; (Roy et al., 2010; Yang et al., 2015)).

Amplification of genomic DNA yielded products for all three primers chosen, while cDNA was only amplified using the primer binding to the −100 bp region relative to the translation start (Figure 1A, gDNA and cDNA, respectively). This confirms the 5’ RACE results by showing that mRNAs containing longer 5’UTRs cannot be detected under the growth conditions, applied within the present study. However, the available annotation of the *NAB1* gene according to Phytozome v12.1 (gene identifier Cre06.g268600) indicated the existence of another alternative transcription start site, at least 361 bp upstream of the translation initiation site. Transcription from two alternative start sites, one of which representing a TATA-box, was reported for other nuclear *C. reinhardtii* genes before (Gromoff et al., 2006; Fischer et al., 2009). Interestingly, a TATA-box and AT-rich region are present in the *NAB1* promoter between −483 to −478 bp and −377 to −358 bp, respectively (Supplementary Table 2) and both could represent sequences of an alternative core promoter.

NAB1 expression is clearly induced under CO_2_-limiting conditions, most notably under mixotrophic cultivation mode based on promoter activation (Berger et al., 2014). In order to narrow down the promoter regions involved in the modulation of *NAB1* promoter activity, reporter constructs were created, in which *Gaussia princeps* luciferase (*g*Luc) expression is driven by a full length (1.55 kb) and different truncated versions of the promoter (Figure 1B). A promoter-less reporter construct served as a control for transformation of *C. reinhardtii* CC-1883 cells (Figure 1B; 0 bp). For each construct, *g*Luc-expressing transformants were found amongst 196 colonies screened (Figure 1B; luciferase expression: +). To examine the carbon dioxide–responsiveness of the distinct promoter versions, three representative transformants were analyzed regarding their differential luciferase expression, under high (3% (v/v)) versus low (0.04% (v/v)) carbon dioxide supply, in acetate-containing medium.

Except for the promoter-less control (0 bp), all other cell lines exhibited a higher luminescence under air levels of CO_2_ compared to carbon dioxide enriched air (Figure 1B; response to CO_2_ limitation: +). This indicated that *g*Luc accumulation following carbon dioxide deprivation requires *NAB1* promoter elements and that even the smallest fragment of 152 bp contains elements conferring CO_2_-responsiveness, albeit the constructs clearly differed concerning their overall induction strength (Supplementary Figure 1).

A time course of *g*Luc accumulation of representative cell lines harboring either the reporter construct containing the minimal promoter (152 bp) or the promoter-less control (0 bp) clearly confirmed that 152 bp are sufficient for promoter activity modulation in response to altered carbon dioxide availability (Figure 1C). While the promoter-less control showed only a moderate response to the withdrawal of carbon dioxide (1.49±0.55 (SEM, n=3) vs. 1.0 under CO_2_-replete conditions at t_4h_), a pronounced induction was seen for the 152 bp fragment (3.86 ± 0.27 (SEM, n=3) vs. 1.0) and the relative difference between the responses of both constructs increased over time.

Due to the fact, that the 152 bp fragment clearly conferred CO_2_-responsiveness to the reporter construct (Figure 1C), we used this sequence as a probe in a Yeast-One-Hybrid experiment (Figure 1D). A *Chlamydomonas* wild-type cell line was grown either under CO_2_-replete conditions or subjected to carbon dioxide depletion. Total RNA was extracted from both cultures and used for the preparation of a cDNA library, with library sequences encoding prey proteins fused to the activation domain of a transcription factor. The 152 bp sequence was cloned upstream of a yeast *His2* gene required for histidine biosynthesis in a histidine auxotrophic strain. Acting as a bait, interaction with the prey within yeast cells enabled transcription of the *His2* gene and thus growth on medium lacking histidine. A total of 34 positive yeast clones were screened and sequenced and among them only clone 14 contained a prey sequence encoding a putative DNA-binding protein (Supplementary data 1). BLAST analyses showed that the prey sequence belongs to a *C. reinhardtii* gene Cre01.g012200.t1.1 (UniProtKB: A0A2K3E5K7), which was named LCRF (Low CO_2_-response factor). This putative DNA-binding protein contains an N-terminal BTB/POZ domain, which was first identified as a conserved motif present in the *Drosophila melanogaster* bric-à-brac, tramtrack and broad complex transcription regulators as well as in many pox virus zinc finger proteins (Zollman et al., 1994; Bardwell and Treisman, 1994; Numoto et al., 1993; Koonin et al., 1992). In combination with zinc finger motifs, BTB domains are frequently implicated in protein-protein interactions leading to homodimerizations of the transcription factor (Ahmad et al., 1998). Besides the BTB domain, the protein contains a highly conserved DNA-binding SBP (squamosa-promoter binding protein) domain, which has a typical size of 79 amino acids and comprises a zinc finger motif with two Zn^2+^-binding sites: Cys-Cys-His-Cys and Cys-Cys-Cys-His (Yamasaki et al., 2004). An isolated SBP-domain is sufficient for the specific recognition of the *cis*-element TNCGTACAA (Cardon et al., 1997; Cardon et al., 1999) and the tetranucleotide GTAC represents the essential core of the motif (Birkenbihl et al., 2005). The 152 bp fragment does not contain a complete GTAC hexanucleotide motif, but it comprises two TAC trinucleotides at positions −79 and −85 relative to the *NAB1* start codon. Presumably, these TAC sequences confer binding to the SBP domain of LCRF in electrophoretic mobility shift assays (Figure 2 A; 152 bp *NAB1* promoter), which can be observed in the presence of excess unspecific competitor (lanes 2-4). Incubation of the recombinant SBP domain with the larger 199 bp probe, however, led to more pronounced complex formation. Importantly, complex formation could be abolished by adding unlabeled specific competitor in excess (199 bp *NAB1* promoter; competitor “+”), thus indicating specificity of the interaction. A small (30 bp) oligonucleotide probe comprising both GTAC motifs also present on the 199 bp probe was analyzed along with a control, containing a shuffled core sequence “CAGT” instead of “GTAC”.

**Figure 2:**
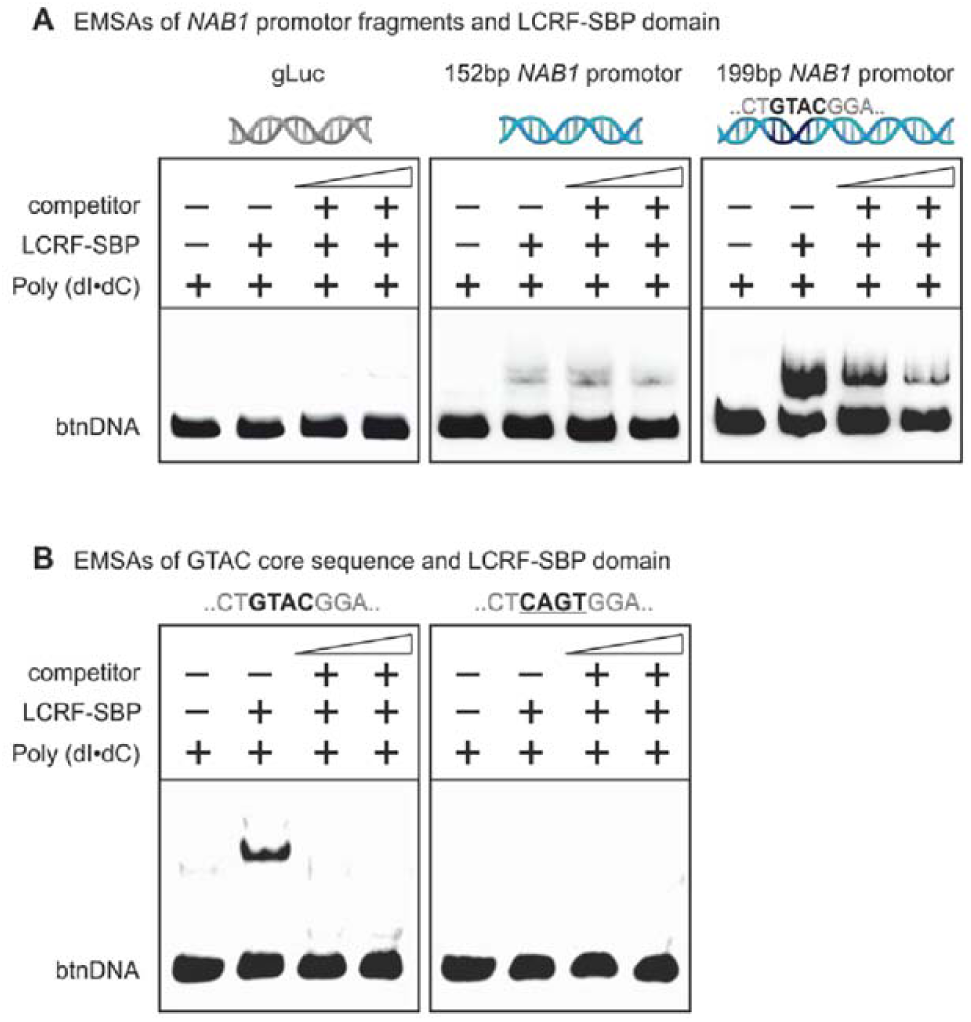
The Squamosa promoter-binding protein (SBP) domain of LCRF binds specifically to a GTAC tetranucleotide motif present in the *NAB1* promoter. Electrophoretic mobility shift assays (EMSA) were performed with recombinant LCRF-SBP and biotinylated DNA probes (btnDNA). **(A)** LCRF-SBP was incubated either with a *gLuc* DNA fragment serving as a negative control, or with two distinct fragments derived from the *NAB1* promoter. All EMSA samples contained the biotinylated probe (*gLuc*, 152 bp or 199 bp fragment) and unspecific Poly (dI:dC) competitor, but differed regarding the presence (“+”) or absence (“-”) of recombinant protein (LCRF-SBP). Biotinylated probes were either added to the protein without (competitor, “-”) or with the simultaneous addition of specific unlabeled competitor (“+”), which was provided in excess (200- and 400-fold relative to the labeled probe) with the highest excess shown in the rightmost lane. **(B)** Biotin-labeled DNA fragments derived from the *NAB1* promoter (position −178 to −149 relative to the 5′ end of the *NAB1* mRNA) either contained intact GTAC cores or cores being disrupted by shuffling (“CAGT”). Probes were incubated with recombinant LCRF-SBP and increasing concentrations (0×, 200× and 400× molar ratio) of unlabeled probe containing the intact GTAC motif (lanes 2–4 of each panel).

LCRF1-SBP bound to the GTAC-containing probe in the presence of excess unspecific competitor (Figure 2C; GTAC), while shuffling of the core (CAGT) abolished binding under these conditions. Complex formation between LCRF1-SBP and the biotinylated, GTAC-containing probe could be outcompeted by adding unlabeled probe. Overall, these *in vitro* binding experiments clearly demonstrated that the SBP domain of LCRF1 specifically binds to the GTAC core motifs present in the *NAB1* promoter at positions −160 and −172 relative to the translation start codon.

### LCRF is essential for the induction of the *NAB1* promoter in response to carbon dioxide limitation

To analyze whether LCRF is indeed required to induce the expression of NAB1 under limiting carbon dioxide conditions, the induction of *LCRF* gene transcription in response to carbon dioxide depletion was first analyzed (Figure 3A; qRT-PCR; wildtype; *LCRF*; light grey bars).

**Figure 3:**
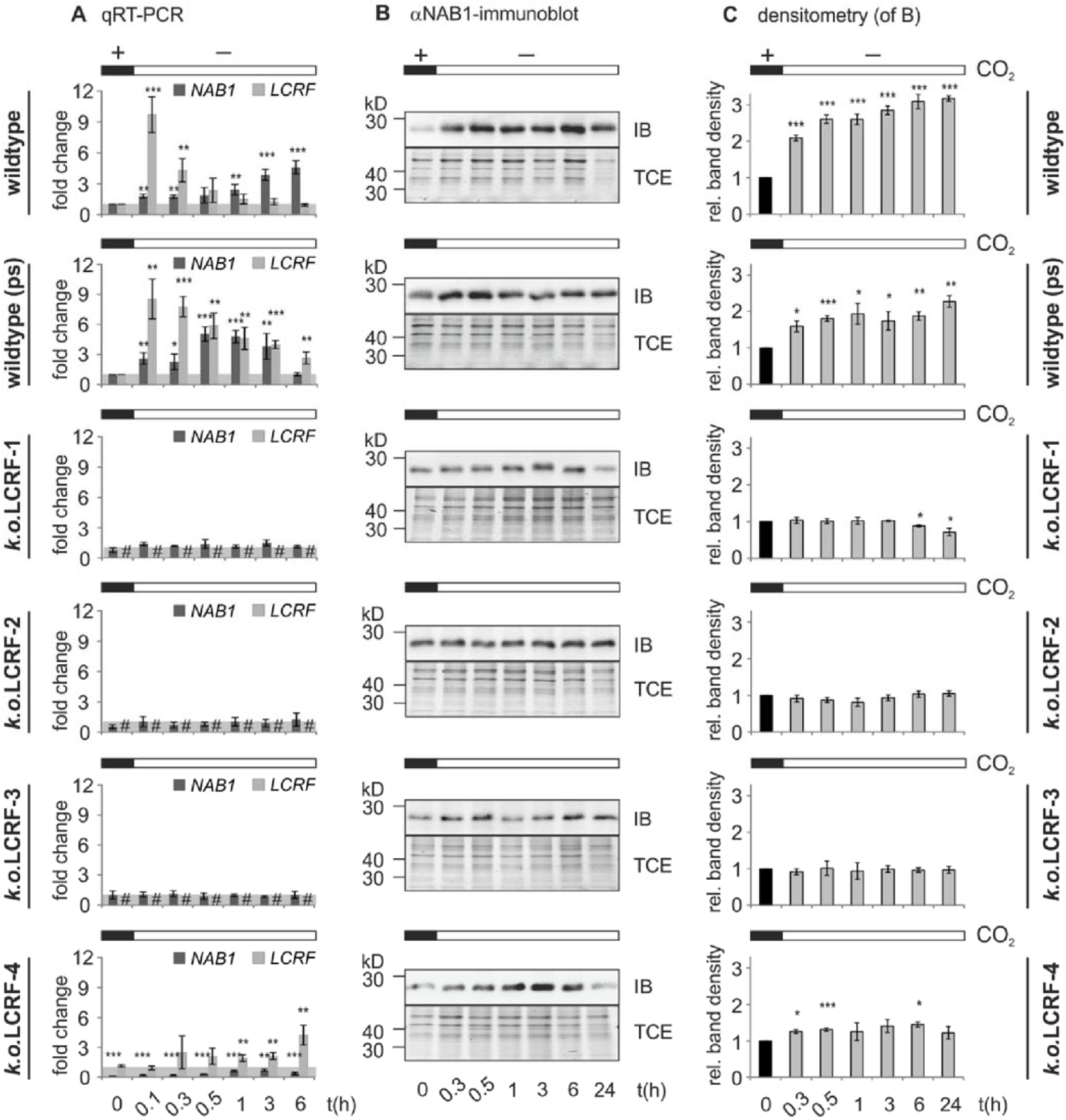
Limited CO_2_ supply activates the expression of the transcription factor LCRF, which in turn enhances the expression and accumulation of NAB1 protein. Cells of a wildtype, the parental strain (wildtype ps) as well as LCRF knock out mutants (*k.o.LCRF* mutants 1 to 4) were cultured photoheterotrophically with high carbon dioxide supply (4% (v/v) CO_2_ in air, time point t_0h_, (+CO_2_)) before subjecting them to low (bubbling with air levels (0.04% (v/v)) of CO_2_) carbon dioxide levels for 0.1, 0.3, 0.5, 1, 3, 6 and 24 h. (**A**) *LCRF* and *NAB1* mRNA levels assessed by qRT-PCR. Wildtype cells, acclimated to 4% (v/v) CO_2_ (t_0h_, (+CO_2_)) served as the reference condition (set to 1). Mean values of three independent experiments are presented. (**B**) Representative images of immunoblot analyses conducted to quantify NAB1 protein levels. Immunoblot signals are shown along with a protein loading control (TCE). (**C**) Densitometric analysis of immunoblot signals, which were normalized to the loading control. Mean values are derived from three independent experiments, each including at least two technical replicates. Error bars represent SD (n = 3 for (A)); SEM (n = 3 for (C)). Asterisks represent p-values as determined via Student’s t-test (* = < 0.05, ** = < 0.01, *** = <0.001) and ‘#’ not detected.

A sudden, hundred-fold drop (4% to 0.04% (v/v) CO_2_ in air; “-”CO_2_) in the availability of carbon dioxide led to a rapid accumulation of transcript LCRF in a *Chlamydomonas* wildtype cell line (9.7±1.7 (SD)-fold at t_0.1h_) within the first 6 minutes after the onset of carbon dioxide deprivation. This steep increase in transcript level was followed by a steady decline, reaching pre-stress levels at about 3 hours after changing the gassing condition (t_0.3h_ to t_3h_). Within this time period *NAB1* transcript levels displayed a steady increase, to reach an about five-fold accumulation (4.6±0.6 (SD)) relative to the carbon dioxide-replete state at the end of the time course (*NAB1*; dark grey bars). The monotonous increase in *NAB1* mRNA levels was correlated with the accumulation of NAB1 protein (Figure 3B and C; wildtype). In response to carbon dioxide limitation, NAB1 protein levels increased about three-fold within 6 hours and remained at this level during the following 18 hours (Figure 3C; hours 6-24). The expression pattern of *LCRF* and *NAB1* transcripts and the time course of NAB1 accumulation further indicated that LCRF is required for NAB1 expression control during low carbon dioxide acclimation. To proof that LCRF is indeed essential for the induction of NAB1 expression under these conditions, we examined four distinct *LCRF* insertion mutants (Supplemental Figure S4) obtained from the *Chlamydomonas* Library Project (CLiP; (Li et al., 2019)) in an identical experimental setup (Figure 3; *k.o.*LCRF1-4). The parental strain of these mutants (Figure 3A; wildtype (ps)) displayed a *LCRF* expression profile resembling the one seen for the *C. reinhardtii* wildtype strain, while *NAB1* transcript levels peaked earlier at around 30 minutes after the onset of carbon dioxide limitation. Carbon dioxide deprivation led to an accumulation of NAB1 protein also in this case, with levels doubling in the course of the experiment (Figure 3C). Importantly, in mutants *k.o.*LCRF1-3 the transcript encoding LCRF could not be detected by qRT-PCR (Figure 3A; *k.o.*LCRF1-3; #) and NAB1 transcript and protein levels remained unchanged following carbon dioxide withdrawal (Figure 3B and C). The fourth mutant showed a largely diminished accumulation of transcript *LCRF* (Figure 3A; *k.o.*LCRF-4), which was accompanied by a lack of NAB1 protein accumulation under low carbon dioxide conditions (Figure 3C). Taken together, these results clearly demonstrate that transcription factor LCRF is essential for the modulation of cellular NAB1 amounts in response to carbon dioxide limitation.

### Inactivation of LCRF leads to an elevated PSII excitation pressure and growth perturbation under carbon dioxide-limited conditions

NAB1 expression analysis in LCRF knock out mutants grown under sufficient vs. limited carbon dioxide supply clearly demonstrated that this transcription factor is crucial for a modulation of cellular NAB1 amounts following an altered inorganic carbon provision (Figure 3). Previous published data clearly showed that an inability to enhance LHCII translation repression via increased NAB1 levels as a response to carbon dioxide limitation results in growth impairment due to elevated PSII excitation pressure (Berger et al., 2014). We therefore analyzed cell division rates, PSII photochemistry and the accumulation of LHCII isoform LHCBM6, as a prime target of NAB1-mediated translation repression (Mussgnug et al., 2005), during the acclimation of parental strain and LCRF mutants to low carbon dioxide levels (Figure 4).

**Figure 4:**
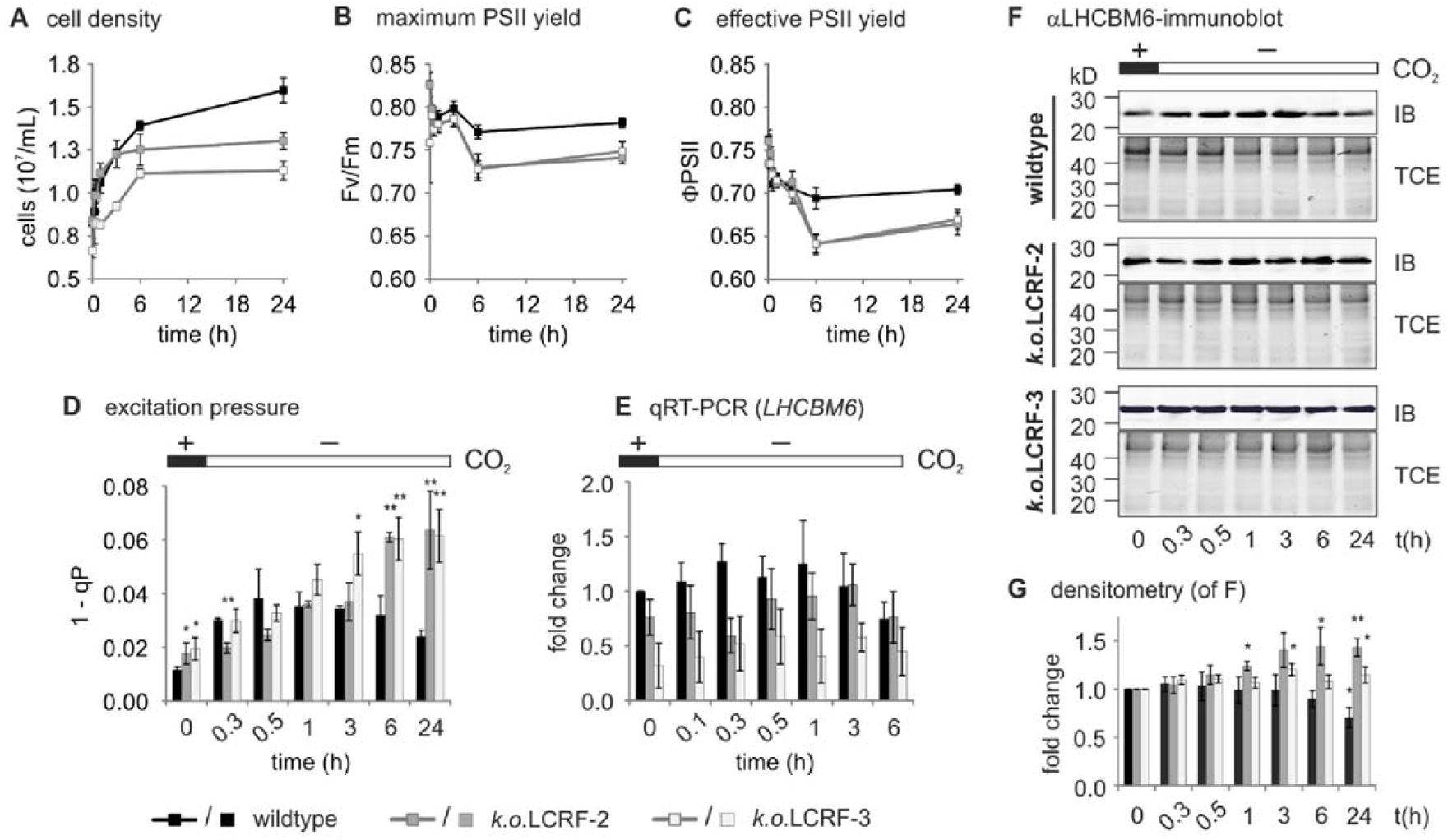
In LCRF knock out mutants, the inability to diminish LHCII protein amounts under limited CO_2_ supply results in an increased PSII excitation pressure and growth perturbation. Parental strain (wildtype) and LCRF knock out mutants (*k.o.*LCRF mutants 2 and 3) were acclimated to high carbon dioxide levels (4 % (v/v); “+”; t_0_) prior to a prompt change of gassing conditions to low levels of carbon dioxide (0.04 % (v/v); “-”; t_0.3h_-t_24h_). Cell numbers (**A**) were determined besides photochemical quenching parameters F_v_/F_m_ (**B**) and ΦPSII (**C**) as well as excitation pressure 1-qP (**D**) at the indicated time points. (**E**) *LHCBM6* mRNA levels as assessed by qRT-PCR. Transcript levels in the wildtype at t_0_ served as the reference condition (set to 1). (**F**) Representative anti-LHCBM6 immunoblot results shown together with a protein loading control (TCE) for wildtype and mutants. (**G**) Densitometric analysis of immunoblot signals, normalized to the loading control presented as mean values derived from three independent experiments (including at least two technical replicates). Error bars represent SD (n = 3 for (E)); SEM n = 3 for (A-D, G)). Asterisks indicate p-values as determined via Student’s t-test (* = < 0.05, ** = < 0.01, *** = <0.001).

Despite starting at almost identical cell numbers at t_0_, prolonged exposure of both LCRF knock out mutants (light and dark grey bars) to air levels of carbon dioxide led to diminished cell numbers (up to 18% and 29% lower for *k.o.*LCRF2 and *k.o.*LCRF3, respectively) compared to the parental strain (black bars) at t_24h_. This suggests that these mutants acclimated less successfully to carbon dioxide limitation than the LCRF-expressing parental strain (Figure 4A; wildtype vs. *k.o.*LCRF2/3). This growth impairment could be further explained by a diminished (≈ 5% lower than parental strain) maximum quantum yield of PSII following exposure to low levels of carbon dioxide in LCRF mutants (Figure 4B; F_v_/F_m_; 6-24h), which indicates a higher susceptibility of PSII to photoinhibition. In addition, PSII photochemistry was altered in the mutants, which could be noted as a reduced photochemical quantum yield of PSII (Figure 4C; ΦPSII; up to 8% lower). The proportion of closed PSII reaction centers (1-qP; Figure 4D) is a measure for the ‘excitation pressure’ of PSII and a continuous rise of 1-qP in the mutants implies an over-reduction of the photosynthetic electron transport chain, which exacerbates during growth in carbon dioxide-limiting conditions. As shown previously (Berger et al., 2014), LHCBM6 transcript levels did not change significantly during the acclimation to low levels of carbon dioxide (Figure 4E). In contrast, prolonged exposure to air levels of carbon dioxide led to diminished LHCBM6 protein levels in the parental strain (Figure 4G; black bars; t_0_ vs. t_6h_ and t_24h_), while the opposite trend could be observed for the mutants (light and dark grey bars). Overall, these data demonstrate that LCRF is a crucial component of the gene regulatory circuit required to adjust NAB1 levels to the prevailing supply of inorganic carbon and that impairment of this circuit affects growth.

## DISCUSSION

In the present study, a novel transcription factor, required for carbon dioxide acclimation in *C. reinhardtii*, was identified. LCRF contains a combination of two conserved domains (Figure 5A; LCRF_Cr), namely, a BTB/POZ (broad-complex, tramtrack, and bric-a-brac/poxvirus and zinc finger) domain at its N-terminus and a C-terminal SBP (Squamosa promoter binding protein) domain (Klein et al., 1996; Zollman et al., 1994; Bardwell and Treisman, 1994; Godt et al., 1993). The BTB/POZ domain is known to facilitate protein-protein interactions (Bardwell and Treisman, 1994; Bonchuk et al., 2011) and can be found in conjunction with many other protein domains (Chaharbakhshi and Jemc, 2016), such as DNA-binding zinc finger domains (Maeda et al., 2005).

**Figure 5:**
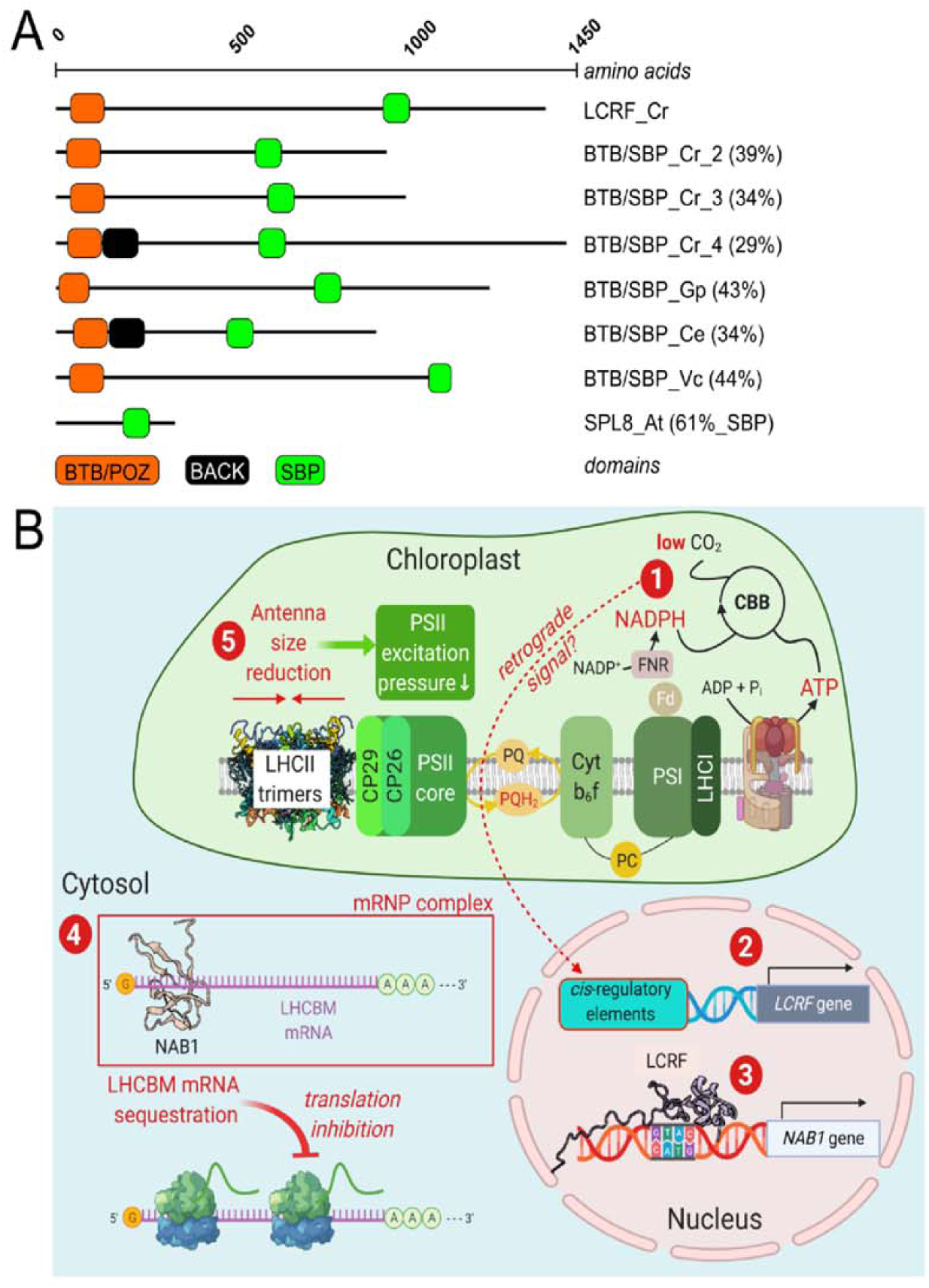
Uniqueness of the BTB/POZ plus SBP combination among known SBP transcription factors and working model of the regulatory circuit for carbon dioxide acclimation based on LCRF action. **(A)** Domain (BTB/POZ, Pfam (http://pfam.xfam.org/) entry PF00651; SBP, Pfam entry PF03110; BACK, Pfam entry PF07707) organization and positions according to the SMART tool (http://smart.embl-heidelberg.de/) of LCRF-homologous proteins identified by BLAST analyses. Amino acid identities (calculated with Clustal Omega; https://www.ebi.ac.uk/Tools/msa/clustalo/) are given in brackets. Abbreviations: Cr: *Chlamydomonas reinhardtii*, Gp: *Gonium pectorale*, Ce: *Chlamydomonas eustigma*, Vc: *Volvox carteri*, At: *Arabidopsis thaliana*. The following amino acid sequences are depicted: LCRF_Cr / UniProt KB accession A0A2K3E5K7; BTB/SBP_Cr2 / A0A2K3CNR5; BTB/SBP_Cr3 / A0A2K3DEV5; BTB/SBP_Cr4 / A0A2K3E2Y5; BTB/SBP_Gp / A0A150FVP1; BTB/SBP_Ce / A0A250XN64; BTB/SBP_Vc / Volvox carteri 2.1 gene identifier Vocar.0002s0008; SPL8_At: Q8GXL3) **(B)** Working model, depicting the regulatory circuit, which adjusts LHCII antenna size to the prevailing supply of inorganic carbon. (1) A limited supply of carbon dioxide slows down the activity of the Calvin-Benson-Bassham cycle and leads to the accumulation of NADPH, which in turn causes an over-reduction of the photosynthetic electron transport chain. (2) A reduced availability of carbon dioxide rapidly activates expression of the LCRF gene via an unknown signaling mechanism, resulting in an accumulation of the *LCRF* transcript. (3) *In vitro* binding data indicate that LCRF binds to GTAC-DNA motifs present in the *NAB1* promoter *in vivo* and activates transcription of the *NAB1* gene. Induction of the *NAB1* promoter eventually leads to elevated cytosolic NAB1 protein levels and enhanced translation repression of *LHCBM*-encoding mRNAs via sequestration in translationally-silent messenger ribonucleoprotein particles (4). A reduced *de novo* synthesis of LHCBM protein reduces PSII antenna size (5) and alleviates PSII excitation pressure.

At its C-terminus, LCRF possesses the plant-specific (Klein et al., 1996; Birkenbihl et al., 2005) DNA-binding SBP domain (Chen et al., 2010), which is sufficient to specifically bind to the GTAC motif present in the *NAB1* promoter (Figure 2B). In total 28 distinct proteins containing an SBP domain are encoded by the nuclear genome of *C. reinhardtii*, as can be seen by BLAST analyses conducted with the SBP domain of LCRF. Apart from LCRF, only one other SBP protein from *Chlamydomonas* has been studied in detail. The copper response regulator CRR1 binds to copper-response elements associated with the *CYC6* promoter that contains a critical GTAC core and drives the expression of a copper-independent substitute for plastocyanin in the photosynthetic electron transfer chain (Kropat et al., 2005). In contrast to *LCRF* (Figure 3), which is induced by carbon dioxide limitation, *crr1* transcription is not activated by copper depletion (Kropat et al., 2005). Copper sensing and CRR1 deactivation occur via binding of Cu^2+^ to the SBP domain, which inhibits DNA-binding (Sommer et al., 2010).

Besides LCRF, three other proteins encoded by the nuclear genome of *C. reinhardtii* combine the BTB/POZ and SBP domain (gene identifiers Cre17.g698233, Cre09.g399289 and Cre02.g110150). Based on the current sequence data deposition available at Pfam 32.0 (El-Gebali et al., 2019), this domain combination seems to be specific for green algal species belonging to the order *Chlamydomonadales* (Fang et al., 2017). This order includes the species *Volvox carteri* (Figure5B; BTB/SBP_Vc), *Gonium pectorale* (BTB/SBP_Vc and UniProt KB accessions A0A150FW26/A0A150GEK1) and *Chlamydomonas eustigma* (BTB/SBP_Ce and A0A250WVT5/A0A250WZD6). The nuclear genome of the higher plant model organism *A. thaliana* encodes 16 SBP-Like (SPL) proteins, which can be subdivided into two families according to their size and sequence similarity (Guo et al., 2008; Xing et al., 2010). The *A. thaliana* SPL most similar to LCRF based on sequence alignment is SPL8, with an identity of 61% within the SBP domain (Figure 5A).

The data shown in the present manuscript allow depicting a chain of molecular events, which enable the fine-tuning of PSII antenna size in response to an altered availability of carbon dioxide (Figure 5B).

A rapid drop in the availability of carbon dioxide slows down carbon fixation in the Calvin-Benson-Bassham cycle, which in turn attenuates the re-oxidation of NADPH, yielding in an over-reduction of the photosynthetic electron transport chain (Lucker and Kramer, 2013) and thus increased PSII excitation pressure (Figure 4D; Figure 5A, event “1”). In a previous study (Berger et al., 2014), we demonstrated that carbon dioxide limitation induces a state transition (Goldschmidt-Clermont and Bassi, 2015) from state I to state II, which starts relaxing after about an hour after the onset of the stress and this transition from state II back to state I continues despite an ongoing carbon dioxide depletion. While the light-harvesting antenna re-associates with PSII, LHCII translation repression via NAB1 is induced to replace the short acclimation strategy based on state transitions by a long-term response, yielding in a PSII antenna, which is adjusted to the prevailing carbon dioxide supply (Berger et al., 2014). NAB1 accumulation in response to carbon dioxide depletion was perturbed in state transition mutant *stt7* (Depège et al., 2003), pointing at a mechanistic link between short and long-term acclimation (Berger et al., 2014). Since an accumulation of NADPH under carbon dioxide limiting conditions also leads to an over-reduction of the plastoquinone pool, as a trigger for state I-to-state II transitions (Bulté et al., 1990; Iwai et al., 2007), this represents a candidate source for a retrograde signal (Chen et al., 2004; Escoubas et al., 1995; Hüner et al., 2012) activating LCRF transcription (Figure 5B; event “2”). Future work will analyse the signalling mechanism underlying LCRF gene induction following carbon dioxide depletion in more detail by applying or instance the *stt7* mutant and inhibitors of the photosynthetic electron transport chain. LCRF transcripts accumulate rapidly, within 6 minutes, after reducing the availability of carbon dioxide (Figure 3A; wildtype). Carbon dioxide depletion is known to activate genes rapidly in *C. reinhardtii*, as was shown for the transcript encoding an inducible mitochondrial carbonic anhydrase, which could be detected already 15 minutes after reducing the carbon dioxide supply from 5% (v/v) to ambient air (Eriksson et al., 1998).

As reported for other SBP-domain containing transcription factors (Birkenbihl et al., 2005), LCRF binds specifically to a GTAC core sequence in the *NAB1* promoter (Figure 2B; Figure 5B, “3”). The *NAB1* promoter contains two of these motifs at positions −160 and −172 relative to the translation start codon. A potential function of the BTB domain could be to facilitate homodimerization or even oligomerisation of LCRF, a mechanism which, in the case of BTB-zinc finger transcription factors, enables cooperative binding to short recognition sequences present in a promoter multiple times (Katsani et al., 1999). Such a mechanism could confer promoter recognition specificity, when binding motifs are small like GTAC and occur frequently in a genome. In addition to be engaged in homodimerization, BTB/POZ domains were also reported to facilitate heterodimerization, thus expanding the spectrum of recognizable motifs via combining distinct DNA recognition specificities (Kobayashi et al., 2000; Stogios et al., 2005). Lastly, mammalian BTB/POZ domains have also been shown to interact with transcriptional repressors such as N-CoR, thereby targeting histone deacetylases to their target genes (Hörlein et al., 1995; Huynh and Bardwell, 1998; Guenther et al., 2001). However, considering that NAB1 accumulation is perturbed in LCRF-free mutants (Figure 3), repressor recruitment by the BTB/POZ domain of LCRF to the *NAB1* promoter seems to be an irrelevant mechanisms for NAB1 expression control.

Although 152 bp were sufficient to mediate carbon dioxide-dependent promoter modulation (Figures 1B and C), longer fragments displayed a stronger induction (Supplemental Figure S1), indicating the presence of additional elements implicated in this response. An *in silico* analysis regarding the presence of known *cis*-regulatory elements on the *NAB1* promoter was performed (Supplementary Figure S2), using the databases PLACE (Higo et al., 1999) and PlantCARE (Lescot et al., 2002). In addition, a recent transcriptome study (Winck et al., 2013) was considered, in which transcription factors and regulators responding to changes in carbon dioxide levels were identified. Besides, the authors could also identify ten sequence motifs and respective motif combinations in promoters regulated by low carbon dioxide. Six of these motifs are present in the *NAB1* promoter (Supplementary Figure S2, Supplementary Table S2), but none of them is located within the 152 bp region upstream of the translation start site. Moreover, according to Winck and co-workers, NAB1 (UniProt KB ID 126810) was contained in a cluster of early responding genes and mRNA levels increased about 7-fold one hour after the onset of carbon dioxide deprivation (Winck et al., 2013).

NAB1 accumulation in response to carbon dioxide limitation requires LCRF (Figure 3C) and results in a decline of LHCBM6 protein levels after six hours (Figure 4F). The transcript encoding LHCBM6 is the prime RNA target of NAB1 (Mussgnug et al., 2005) and previous work has already demonstrated that enhanced LHCBM6 translation repression, based on elevated NAB1 levels under carbon dioxide limitation (Figure 5B; “4”), leads to a strong PSII antenna size reduction of about 50% (Berger et al., 2014). This antenna size reduction is essential for maintaining a low excitation pressure despite an ongoing carbon dioxide limitation (Berger et al., 2014) and a lack of antenna size control in LCRF mutants results in a high excitation pressure after prolonged exposure to carbon dioxide depletion (Figure 4D). NAB1-mediated translation repression therefore prevents a rise in PSII excitation pressure (Figure 5B; “5”), when carbon dioxide limitation lasts for several hours. Overall, the data presented allow a detailed depiction of the regulatory circuit (Figure 5B), which is required to adjust the algal light-harvesting capacity to the prevailing supply of inorganic carbon. This circuit implicates the newly identified transcription LCRF as well as the translation repressor NAB1 (Mussgnug et al., 2005; Wobbe et al., 2009; Berger et al., 2014; Berger et al., 2016; Blifernez et al., 2011) and will contribute to an improved understanding of long-term carbon dioxide acclimation in microalgae.

## METHODS

### Strains and culture conditions

Wild-type *C. reinhardtii* strain CC-1883 (cw15 mt-; Chlamydomonas resource center, St. Paul, MN, USA) and derived reporter strains (see below, Supplementary Figure S1) were grown with acetate (tris-acetate-phosphate media; (Harris et al., 2009)), in continuous white light at 250 μmol photons m^-2^ s^-1^ and bubbled with air or carbon dioxide enriched (3% (v/v)) air, as described before (Berger et al., 2014). For the analyses concerning the LCRF mutants (LMJ.RY0402.207557_2, LMY.RY0402.14.1786_1, LMJ.RY0402.121192_1, LMY.RY0402.246869_1; referred as *k.o.*LCRF 1, 2, 3 and 4 mutants (Supplementary Table S3; (Li et al., 2019)) as well as the parental (CC-4533, cw15 mt- [Jonikas CMJ030; (Zhang et al., 2014)], Chlamydomonas resource center) and wildtype (CC-1883) strains, the cultivation was performed as follows: Photoheterotrophically cultured (pre-adapted to 4% (v/v) CO_2_ in air and 400 µmol photons m^-2^s^-1^ continuous white light, time point t=0 h, (+CO_2_)) cells were subjected to low carbon dioxide levels (0.04% (v/v) by bubbling the cultures with air (–CO_2_) for 0.1, 0.3, 0.5, 1, 3, 6 and 24 h. The validation of the mutant strains (Supplementary Figure S4, Supplementary Table S3) was performed following the “Instructions for characterizing insertion sites by PCR” available on https://www.chlamylibrary.org, using the primers listed in Supplementary Table S1.

### Fluorescence analyses

Chlorophyll fluorescence was measured for the wildtype and *k.o.*LCRF mutants in parallel in four technical replicates per measurement for each time point in a 96 well plate (flat bottom) using a closed FluorCam FC 800-C Video Imager (Photon Systems Instruments, Brno, Czech Republic). Before the measurement, samples were dark adapted for at least 1 h. The evaluation of the efficiency of PSII photochemistry as well as photochemical quenching parameters was calculated as described (Maxwell and Johnson, 2000; Murchie and Lawson, 2013).

### Determination of the transcription start site of gene NAB1

A 5’RACE analysis was performed with modifications according to Scotto-Lavino et al. to identify the length of the 5’UTR in the *NAB1* gene (Scotto-Lavino et al., 2006). Total RNA was extracted from strain CC-1883 after cell harvesting in the late-logarithmic phase of a photoheterotrophic cultivation and purified as described below. Uncapped RNA was excluded from adapter ligation by using alkaline phosphatase and tobacco acid pyrophosphatase. The residual RNA, regarded as complete, was converted into cDNA and amplified using two nested primer pairs (Supplementary Table S1; 5’RACE). PCR products were sequenced (MPIZ DNA core facility on Applied Biosystems; Weiterstadt, Germany). The results of the 5’RACE were validated by performing PCRs (Supplementary Table S1; mapping of TSS) on gDNA and cDNA of strain CC-1883. Products appearing for both were considered to be formed within the transcribed region of NAB1 in contrast to those only formed with gDNA as template.

### Experimental promoter analysis - Vector construction, strain generation and reporter assay

Three strategies were applied to obtain vectors with truncated *NAB1* promoters. For the 817 and 521bp fragments, the *g*Luc expression vector containing a 1.55 bp *NAB1* promoter fragment (Berger et al., 2014) was cut with the FastDigest^®^ (Thermo Scientific) restriction endonuclease SpeI (cutting two times) or *Xba*I and *Avr*II (cutting one time each). This removed a 5’upstream region, whereas an 817 bp or 521 bp fragment, respectively, relative to translation start remained. After purification, subsequent self-ligation let to the construction of pNabCAgLuc_817 and pNabCAgLuc_521. Fragments containing 398, 283 or 152 bp upstream of translation start were amplified (primers are listed in Supplementary Table S1), cut with *Xba*I and *Nde*I and ligated into the vector backbone. Third, an oligo sequence obtained from annealing Pnab_del_fw and Pnab_del_rv (Supplementary Table S1) was inserted instead of a promoter fragment as a negative control. All vectors were checked by sequencing (MPIZ DNA core facility on Applied Biosystems; Weiterstadt, Germany).

Transformation of CC-1883, screening and luminescence assay was performed as described before (Berger et al., 2014). Three independent cell lines per construct were investigated in at least two biological and three technical replicates (except for the 398 bp fragment where only two cell lines were examined). For all strains analyzed, the insertion of the *NAB1*::*g*Luc fragment was verified by PCR.

### Yeast-One-Hybrid experiment

The *C. reinhardtii* wildtype strain CC-1883 was grown with 3% (v/v) CO_2_ and an illumination of 100 µmol photons m^-2^s^-1^ in TAP medium until mid-log phase. A sample (“+CO_2_” condition) for RNA isolation according to the method of Chomczynski & Sacchi (Chomczynski and Sacchi, 1987) was taken and the gassing condition switched to bubbling with air (0.04% (v/v) CO_2_). After 4 h of cultivation carbon dioxide-deplete conditions, RNA was isolated for the “–CO_2_” condition. Pooled RNA samples derived from each conditions were subjected to DNaseI digest (RNase-free DNaseI, Promega) in the presence of RNase inhibitor (RNasin Plus, Promega) according to the manufacturer’s instructions. Dnase digested RNA was further purified using an RNA Clean & Concentrator™-25 (ZYMO RESEARCH) according to manufacturer’s instructions. Purified RNA was used for cDNA library preparation and Yeast-One Hybrid screening at Creative Biolabs Inc. For Y1H screening, the 152 bp *NAB1* promoter fragment upstream of the translation start site was used as bait. The construction of the yeast one-hybrid bait vector pHIS2 and the screening of the yeast cDNA single-hybrid library (pGADT7 Library) the CLONTECH Yeast One-Hybrid System was used. A total of 34 positive clones were screened and sequenced. The obtained sequences were analyzed using the databases NCBI, Phytozome (https://phytozome.jgi.doe.gov; (Goodstein et al., 2012), PlantTFB (http://planttfdb.cbi.pku.edu.cn; (Jin et al., 2017; Tian et al., 2020), Plant Transcription Database, plntfdb (http://plntfdb.bio.uni-potsdam.de/v3.0/; (Pérez-Rodríguez et al., 2010). Except for ten clones, the remaining 24 clones were 22 different protein-coding genes (Supplementary Data 1).

### In silico promoter analysis

The databases PLACE (http://www.dna.affrc.go.jp/PLACE; Higo et al., 1999) and PlantCARE (http://bioinformatics.psb.ugent.be/webtools/plantcare/html; Lescot et al., 2002) were used to search for known *cis*-regulatory element within the 1548 bp fragment upstream of the NAB1 translation start site. Furthermore, motifs which are connected to CO_2_-responsiveness as identified in a transcriptome study (Winck et al., 2013), were taken into account. Additionally, PlantRegMap (http://plantregmap.cbi.pku.edu.cn/; (Jin et al., 2017)) was used to screen the *NAB1* promoter for putative transcription factor binding sites. All *in silico*-identified elements within the *NAB1* promoter are listed in Supplementary Table S2 and Supplementary Figures S2 and S3.

### Electrophoretic mobility shift assays (EMSA)

The interaction of the *NAB1* promoter fragments with recombinant LCRF-SBP (rLCRF-SBP) was analyzed by using the LightShift^®^Chemiluminescent EMSA Kit (Thermo Scientific™), following the manufacturer’s instructions. The 152 bp and 199 bp *NAB1* promoter fragment upstream of the translation start site was amplified using primers Pnab-152_btn_fw, Bnab-199_btn_fwd and Pnab0_rv (Supplementary Table S1). As a control for unspecific protein binding, a similarly sized fragment of the g*LUC* coding region was amplified using gLuc+164_btn_fw and gLuc+316_rv. The forward primers integrate a biotin-TEG-tag at the 5’end of the respective PCR product. Additionally, the same fragments were amplified by using unlabeled forward primer und served as specific unlabeled competitor. The resulting PCR fragments (Supplementary Figure S5B) were purified by applying the ExoSAP-IT™ PCR Product Cleanup (Thermo Fisher Scientific, Affymetrix Inc.) according to the manufacturer’s instructions. In addition, short 30-bp DNA (non-)biotinylated oligos derived from the *NAB1* promoter (position −178 to −149 relative to the 5′ end of the *NAB1* mRNA) with either intact GTAC cores or with cores being modified by shuffling (“CAGT”) (GTAC-consensus and mutated GTAC-consensus sequences, respectively; Supplementary Table S1). Appropriate oligonucleotide pairs were combined in annealing buffer (10 mM Tris-Cl pH .5, 1 mM EDTA, and 10 mM MgCl_2_) to yield a concentration of 50 μM for each oligonucleotide. After boiling for 10 min, the sample was slowly cooled down to 20°C with a cooling rate of 0.2 °C per min.

### Cloning, Expression, and Purification of rLCRF-SBP protein

A 501 bp long coding sequence (including the SBP domain) derived from the LCRF-protein (Cre01.g012200, amino acid number 866-1032, NCBI Accession number: PNW88054) was codon optimized for *E. coli* and *de novo* synthesized (Genscript) including a five amino acid long C-terminal linker sequence and an eight amino acid long His-tag. Cloning was performed using the restriction enzymes *BamH*I and *Xho*I and ligation into the vector pET24a(+) (Novagen). Transformation of chemically-competent *E. coli* KRX cells (Promega) was performed via heat shock method and subsequent selection on LB plates containing 50 mg/L kanamycin. Protein production was induced by addition of 0.1% (w/v) rhamnose at OD_600_ ∼ 0.4 and subsequent cultivation overnight at 24°C. Cells were harvested via centrifugation (5,000 xg, 10 min, 4°C), resuspended in cold PBS including cOmplete™ Protease Inhibitor Cocktail (Merck) and lysed via sonication. After centrifugation (18,000 xg, 10 min, 4°C) recombinant SBP protein (rSBP) was purified via Ni-NTA affinity chromatography. Corresponding protein samples were separated during SDS-PAGE prior to colloidal Coomassie staining (Dyballa and Metzger, 2009). Purified rLCRF-SBP (eluted fraction No. 3, Supplementary Figure S5A) was used for electrophoretic mobility shift assays (EMSAs).

### RNA isolation and quantitative real-time RT-PCR

Total *C. reinhardtii* RNA for qRT-PCR analyses was isolated by using *Quick*-RNA™MiniprepKit (Zymo Research) according to manufacturer’s instructions. Quantitative real-time RT–PCR (qRT–PCR) was carried out using the SensiFast™SYBR Hi-ROX One-Step Kit (Bioline) as described (Wobbe et al., 2009). *NAB1* and *LCRF* mRNA was amplified by use of oligonucleotides Nab1-qpcr4_lp/ Nab1-qpcr2_rp and LCRF-rtq_fwd/rev, respectively (Supplementary Table S1). The determination of the expression of *LHCBM6* and *ACTIN* mRNA was performed as described previously (For sequence details see, (Wobbe et al., 2009).

### SDS-PAGE, immunoblotting and Densitometrical Scanning

Prior to immunoblotting, proteins were separated by 12% Tris–Tricine or Tris–Glycine-SDS-PAGE, with the resolving gel containing 2% (v/v) 2,2,2-Trichloroethanol ((TCE), method adapted from Ladner et al. (Ladner et al., 2004)). Protein amount was determined by application of the Lowry assay (DC Protein Assay, Bio-Rad, CA, USA). After separation, the proteins were visualised using UV light (300 nm), as this causes a photoreaction of TCE with the tryptophan residues of the protein’s side chain (Ladner et al., 2006). Immediately, after TCE in-gel visualization (served as loading control) the same gel was electroblotted to a PVDF membrane (0.2 µm) and the immunodetection was performed using enhanced chemiluminescence (ECL; GE Healthcare). Anti-NAB1 antiserum was generated as described (Mussgnug et al., 2005) and anti-LHCBM6/8 (formerly LHCBM4/6) was a kind gift of M. Hippler (Münster, Germany). This antibody recognizes two distinct LHCBM isoforms, namely LHCBM6 and LHCBM8. For densitometric quantification, the software GelAnalyzer 2010a (Lazarsoftware, Hungary) was applied.

### Statistics

Statistical analysis was performed with two-tailed Student’s t-test, resulting in p-values indicated by asterisks (p≤0.05 = *, p≤0.01 = **, p≤0.001 = ***). Results were shown either, as mean value, or fold-change of mean values. Error bars represent standard deviation or standard error of mean (SD/SEM).

## AUTHOR CONTRUBUTIONS

O.B.K., H.B., V.K., L.W. and O.K. conceived this study. O.B.K., H.B., L.W. and O.K. supervised experiments and analyses. H.B. and O.B.K. performed all *in silico* NAB1-promoter analyses. H.B. analyzed the 5’UTR of the *NAB1* gene. H.B., L.S. and T.Bu. conducted all promoter truncation experiments as well as the expression analyses of the promoter-reporter constructs. M.S. performed sample preparation for the cDNA library for Y-1-H-Experiments. O.B.K., T.Ba. and B.M. performed the EMSA studies. O.B.K., V.K., and B.M. accomplished all studies concerning the analyses of *k.o.*LCRF-mutants. O.B.K., H.B., L.W. and O.K. wrote the original draft manuscript with contributions from all authors. O.B.K., L.W. and O.K. reviewed and edited the manuscript. All authors read and approved the final manuscript.

## ACKNOWLEDGEMENTS

The authors would like to acknowledge the Deutsche Forschungsgemeinschaft (KR 1586/10-1) for funding. The authors would like to thank M. Hippler for providing the antibody against LHCBM6/8. We are grateful to the Center for Biotechnology (CeBiTec) at Bielefeld University for access to the Technology Platforms. No conflict of interest declared.

## Ethics approval and consent to participate

Not applicable.

## SUPPLEMENTAL MATERIALS

Supplemental Information and a Supplementary Data Set 1 are available as separate files.

## SUPPLEMENTAL INFORMATION

**Figure S1.**
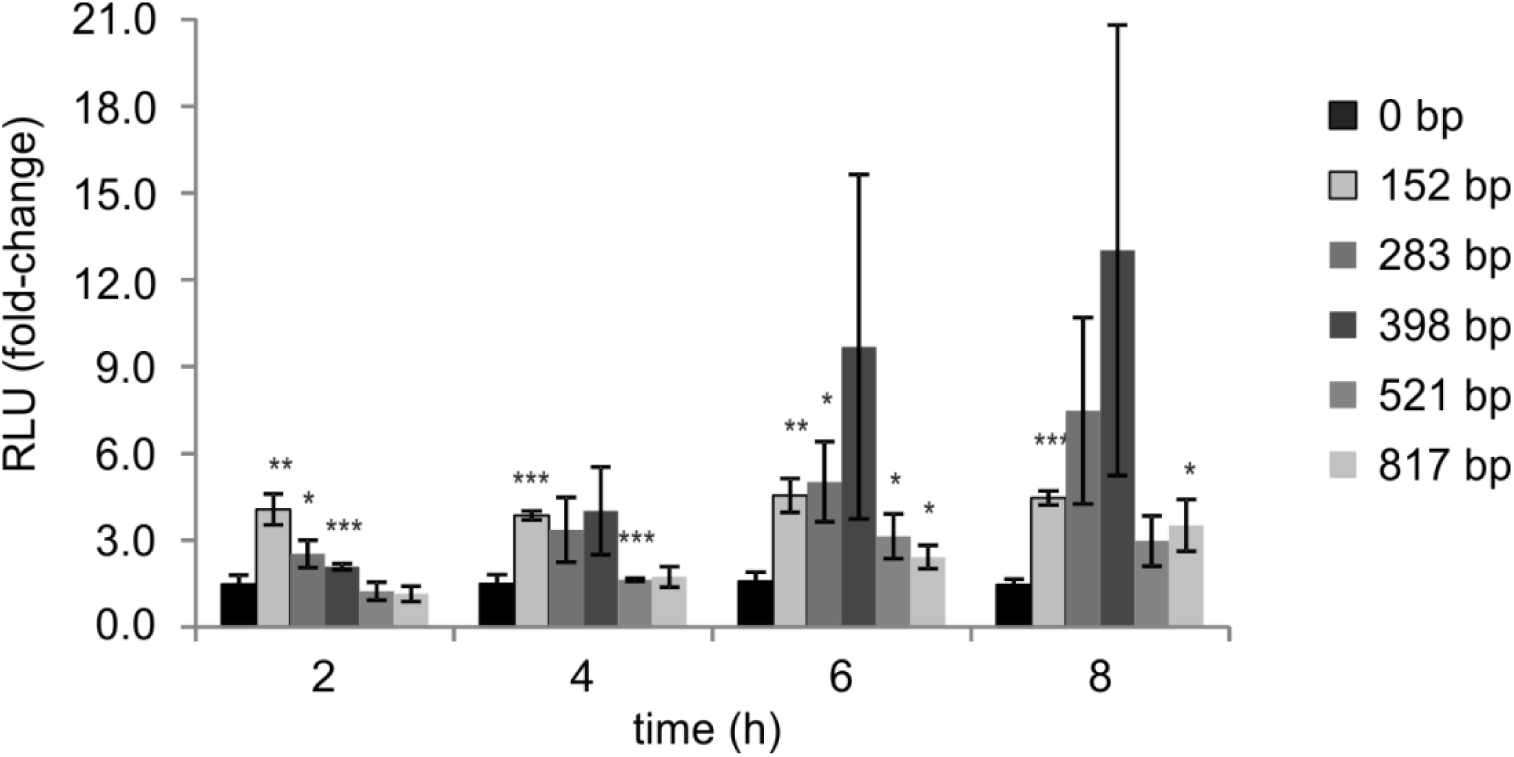
Expression induction analysis of CO2-deprived transformants harboring truncated *NAB1* promoter fragments, (related to Figure 1) Illustrated are the luminescence assay results of carbon dioxide deprived transformants containing a stably integrated gLuc reporter either driven by a *NAB1*-promoter fragment sequence (152 bp, 283 bp, 398 bp, 521 bp, 817 bp) or being devoid of a promoter (0 bp control). For each construct, the luminescence determined under CO_2_-replete conditions (3% (v/v) CO_2_) at t0 was set to 1. Error bars represent the standard error derived from experiments using three distinct cell lines per each construct (with the exception of the 398 bp fragment where only two cell lines were examined) and include the mean values of at least two biological and three technical replicates per cell line (SEM, n=2 for 398 bp-fragment; n=3 for all other fragments). Asterisks represent p-values as determined via Student’s t-test (* ≤ 0.05, ** ≤ 0.01, *** ≤ 0.001).

**Figure S2.**
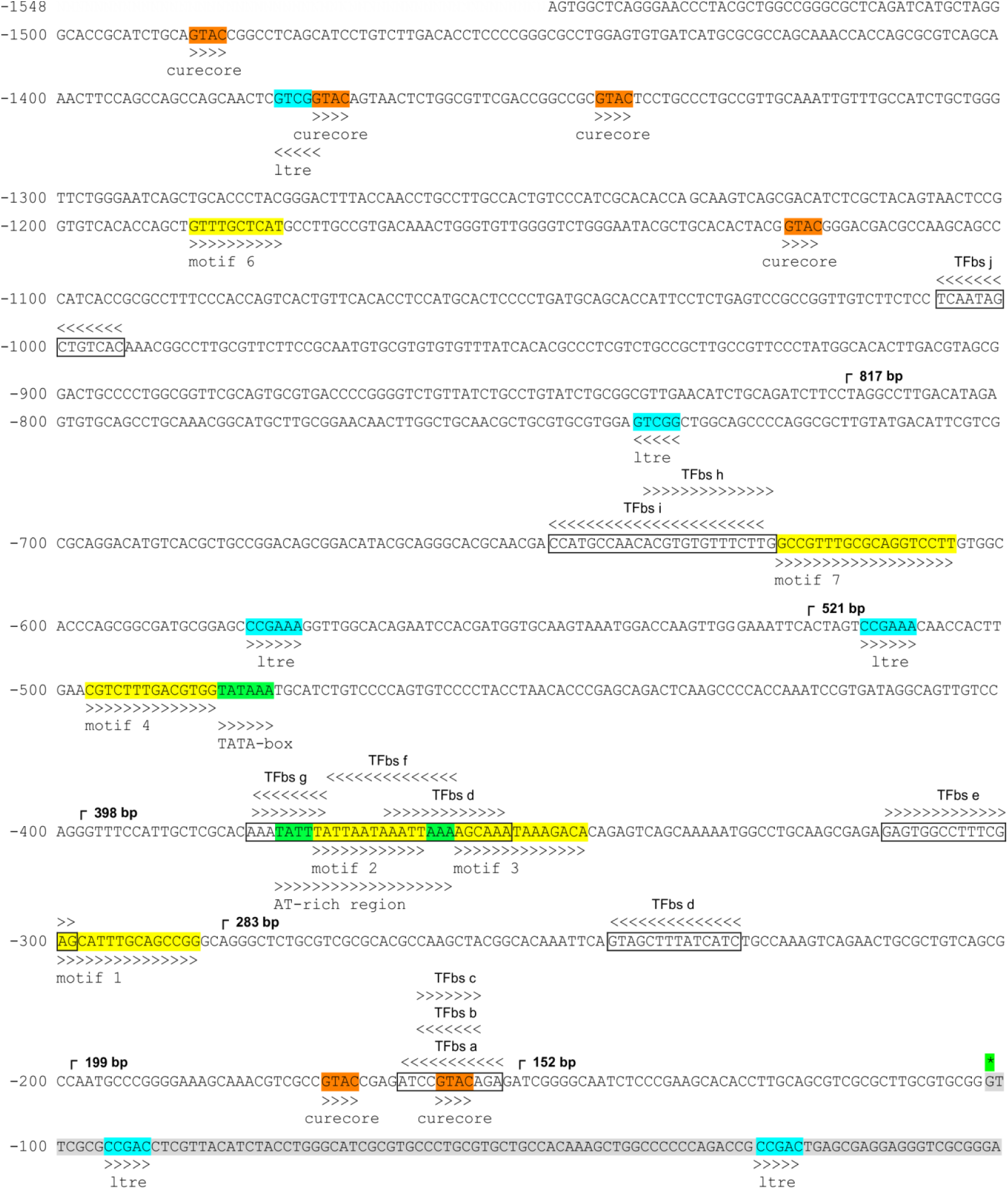
Detailed annotation of *NAB1* promoter sequence, (related to Figures 1 and 2) Illustrated are the candidate *cis*-regulatory elements (CREs)^1^ and transcription factor (TF)^2^ binding sites in the 1548 bp element upstream of *NAB1* translation start. For details see Table S2 and Supplementary Notes.

**Figure S3.**
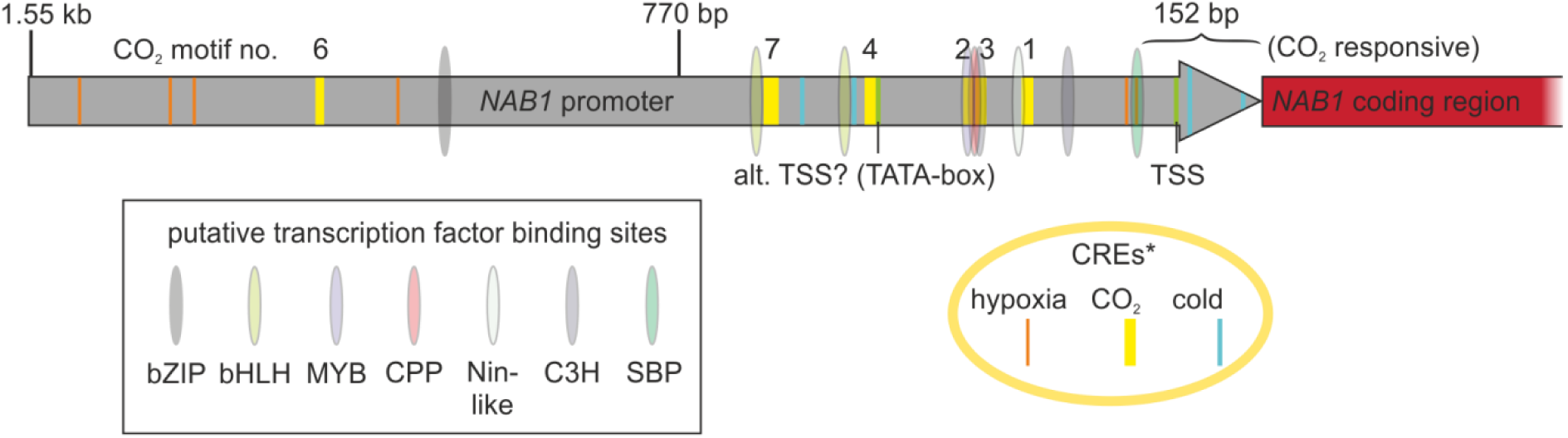
Summarized view on the *NAB1* promoter encoding candidate CREs^1^ and putative transcription factor binding sites^2^, (related to Figures 1 and 2) For details see Figure S2, Table S2 and Supplementary Notes.

**Figure S4.**
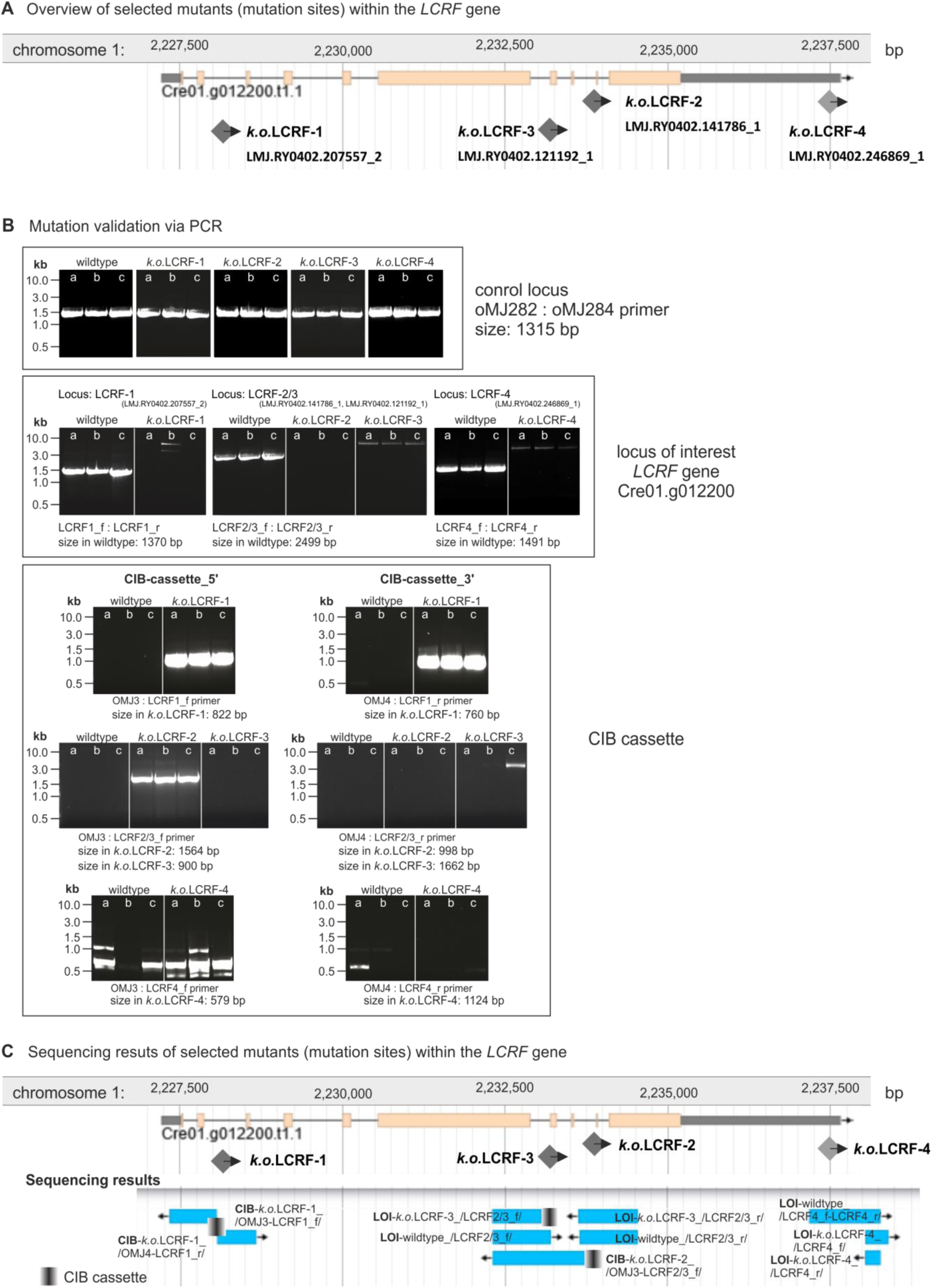
Validation of the LCRF mutants via PCR amplification, (related to Figure 3) The validation of the mutant strains was performed following the “Instructions for characterizing insertion sites by PCR” available on https://www.chlamylibrary.org.

**Figure S5.**
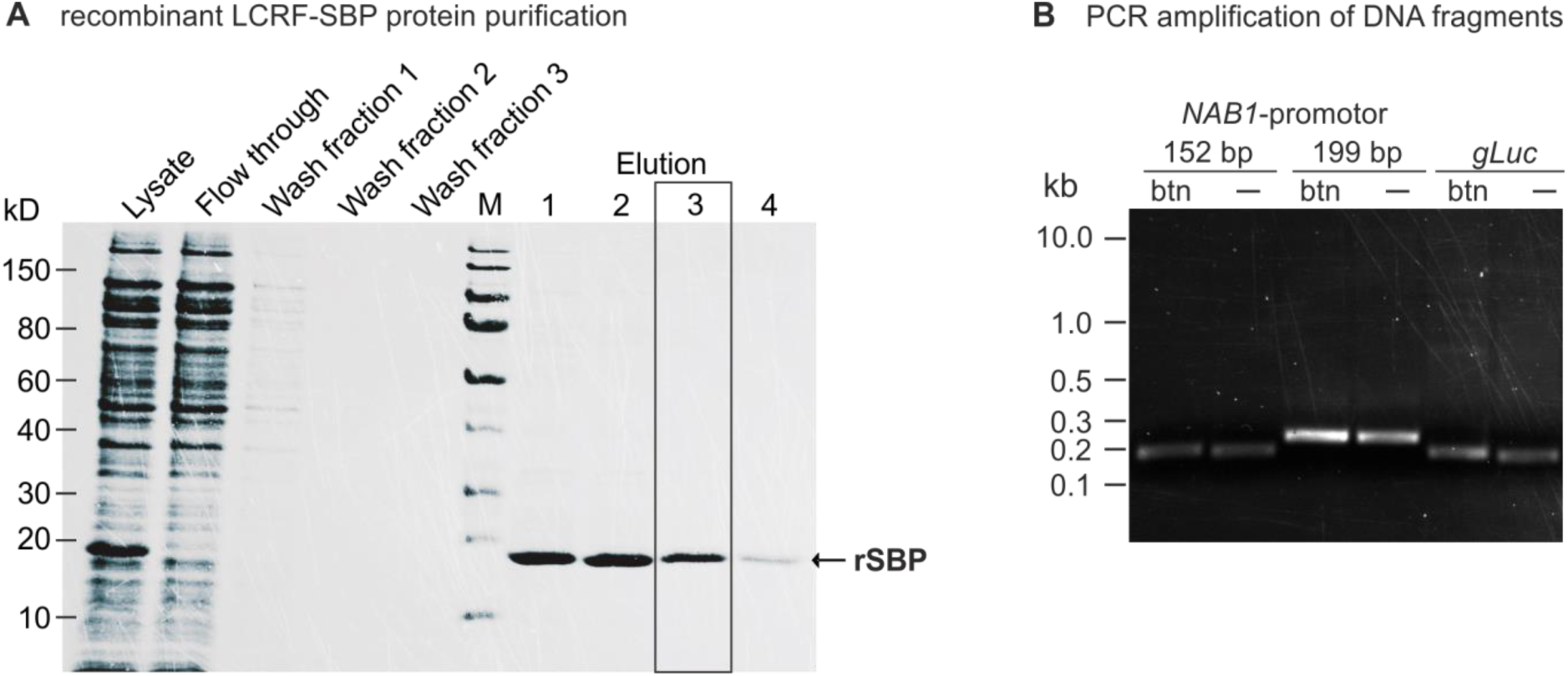
Generation of purified rLCRF-SBP protein and DNA probes for EMSA, (related to Figure 2) Illustrated is (**A**) the purification procedure outcome of the recombinant LCRF-SBP protein and (**B**) the amplified 152 bp and 199 bp *NAB1* promoter fragment upstream of the translation start site as well as *gLuc* coding region (control for unspecific protein binding). The DNA probes were amplified by using unlabeled as well as biotin-TEG-tagged (at the 5’end) forward primer.

**Table S1.**
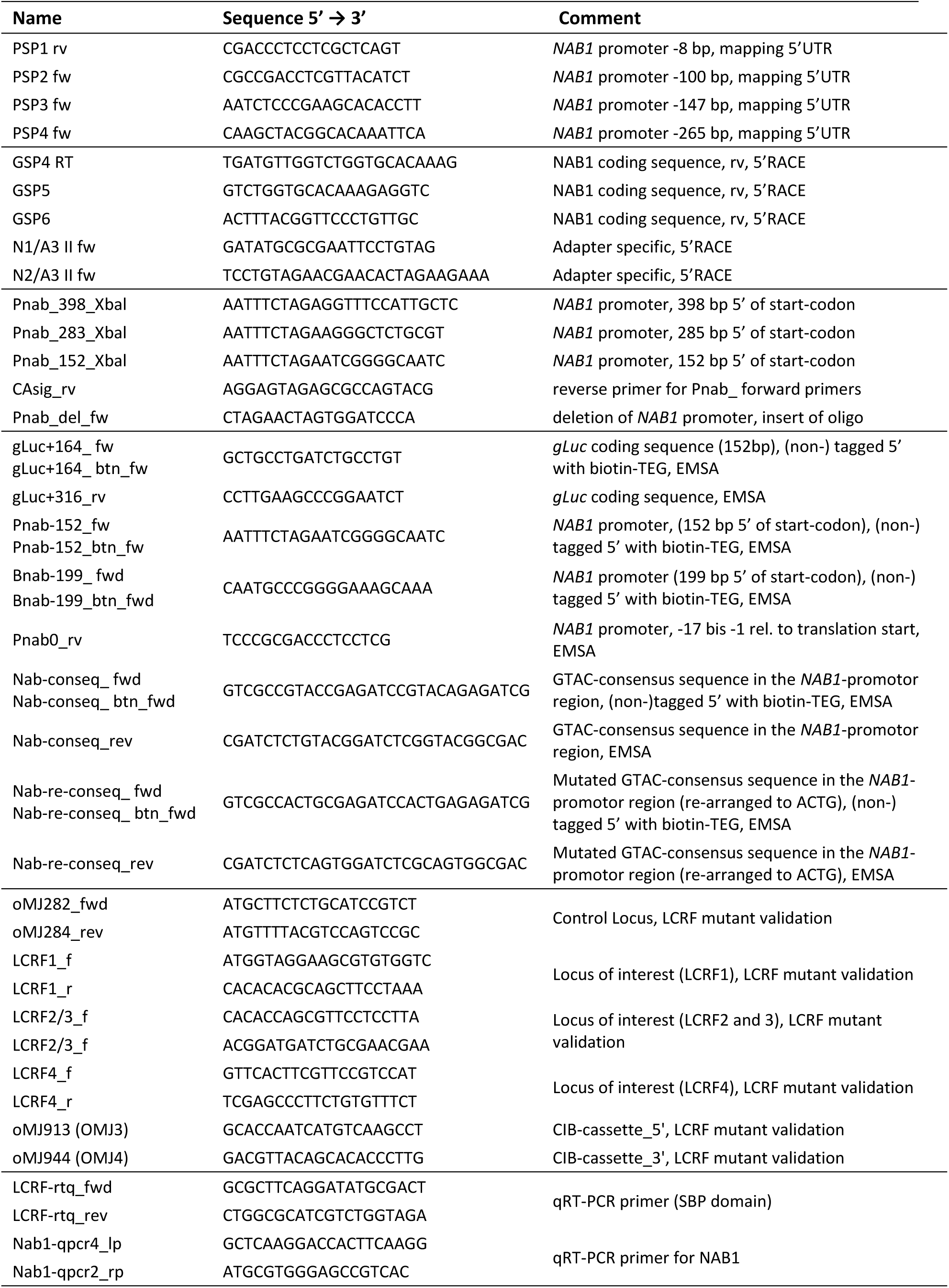
DNA oligonucleotide sequences, (related to Figures 1, 2, 3 and 4).

**Table S2.**
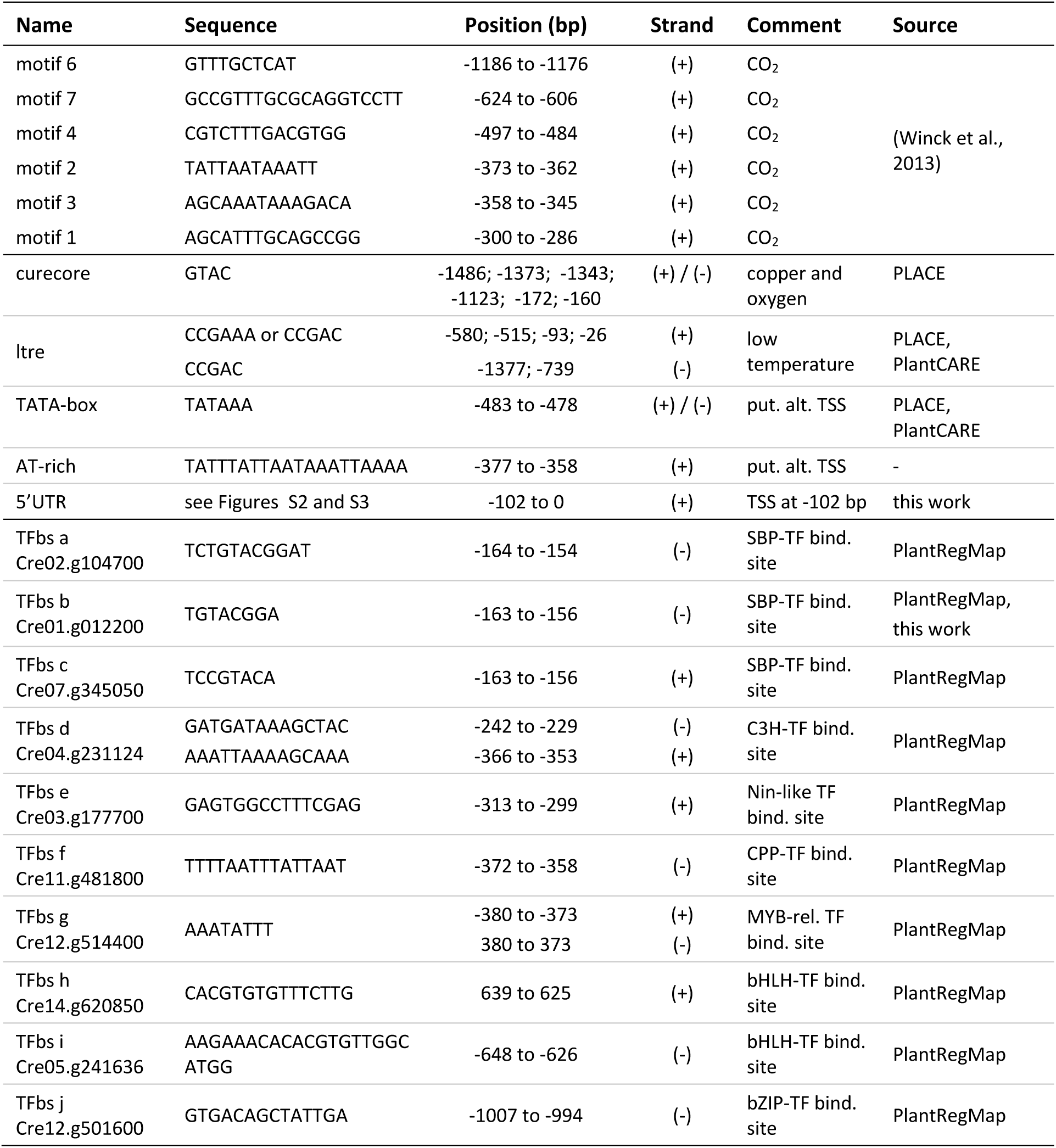
*In silico* identified putative motifs, CREs and transcription factor binding sites within the *NAB1* promotor (1548 bp upstream of the translation start), (related to Figures 1, 2 and 5) Description: CO_2_, copper and oxygen, low temp. (temperature): element confers responsiveness to respective factor; put. alt. TSS: core promoter sequence of putative alternative transcription start; CREs: cis-regulatory elements; TF bind. site (TFbs): Transcription factor binding site; (+) current strand; (-) opposite strand Used databases PLACE (http://www.dna.affrc.go.jp/PLACE; (Higo et al., 1999)) and PlantCARE (http://bioinformatics.psb.ugent.be/webtools/plantcare/html; (Lescot, 2002)), PlantTFB (http://planttfdb.cbi.pku.edu.cn; and http://plantregmap.cbi.pku.edu.cn/ (Jin et al., 2017; Tian et al., 2020)

**Table S3.**
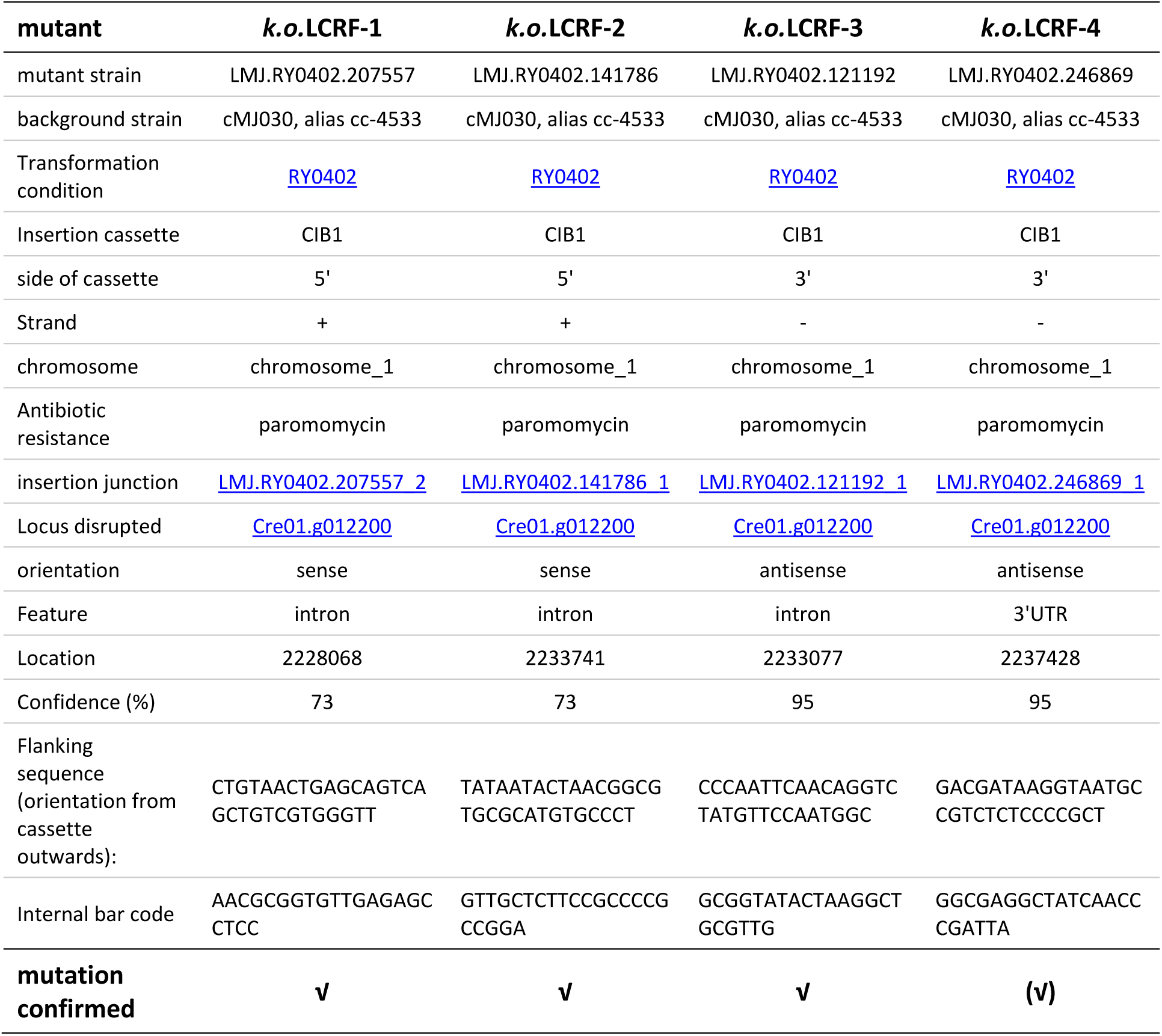
Characteristics of the LCRF mutants, (related to Figures 3 and 4)

## Supplementary Notes

### Chapter 1 *In silico* analyses of putative motifs and CREs in the *NAB1* promotor sequence

To gain further insight on the location of *cis*-regulatory elements within the promoter, the 1.55 kb element was analyzed in silico regarding the presence of known *cis*-regulatory elements applying the databases PLACE (http://www.dna.affrc.go.jp/PLACE; (Higo et al., 1999)) and PlantCARE (http://bioinformatics.psb.ugent.be/webtools/plantcare/html; (Lescot, 2002)) as well as a recent transcriptome study (Winck et al., 2013). In the latter publication, transcription factors and regulators which are controlled by the availability of carbon dioxide were identified and the authors could narrow down ten sequence motifs and respective motif combinations in promoter regions regulated by low carbon dioxide. Intriguingly, six of these motifs are present in the NAB1 promoter (Table S2), however none of them within 152 bp upstream the translation start site (Figure S2). Nevertheless, this study confirms the induction of *NAB1* transcription under CO2 limitation. *NAB1* was grouped into a cluster of early responding genes, and mRNA levels were increased by factor 6.9 after one hour, 11.1 after two hours and 3.2 after three hours (Table S3 of (Winck et al., 2013)).

The databases PLACE and PlantCARE focus on regulatory elements identified in vascular plants, but also motifs of *C. reinhardtii* were added. Using these tools, numerous elements can be found on the *NAB1* promoter, and the most relevant, as they were originally detected in *C. reinhardtii* and/or are associated to carbon metabolisms or light-harvesting regulation, are described here (Figure S2 and summarized in Figure S3).

Four elements responding to low temperatures, but which are also involved in light signaling, are present in the *NAB1* promoter (Table S2, ltre); the sequence CCGAAA originally described in in barley (Dunn et al., 1998) at position −580 bp and −515 bp, and CCGAC identified in *A. thaliana* (Kim et al., 2002) and winter *Brassica napus* (Jiang et al., 1996) within the *NAB1* 5’UTR at −93 bp and −26 bp before translation start site. Furthermore, an element that confers responsiveness to copper and oxygen deficiency in *C. reinhardtii* (Quinn et al., 2002; Kropat et al., 2005) is encoded six times on *NAB1* promoter (Table S2, curecore).

An enhanced expression of NAB1 during cold periods and hypoxia seem reasonable. Low temperatures slow down metabolic reactions such as the Calvin cycle, and oxygen limitation decreases the consumption of reducing equivalents in the mitochondrial electron transport chain. Both causes an over-reduction of the photosynthetic electron transport chain and increases photosystem II excitation pressure. NAB1 mediated repression of LHCBM protein synthesis could lower the PSII antenna size under cold or hypoxic conditions and relieve the pressure.

Two enhancer elements with the consensus motif GANTTNC, the binding sites site of the transcription factor LCR1 to the *CAH1* promoter, are crucial for expression induction of the carbonic anhydrase 1 under carbon dioxide limitation (Kucho et al., 2003; Yoshioka et al., 2004). The consensus sequence is found five times on the *NAB1* promoter, but exclusively on reverse strand, indicating that NAB1 is might not be regulated via the same pathway as the factors involved in the carbon concentrating mechanism.

Also light responsive elements can be detected on the *NAB1* promoter; for instance the GATA-box, which is conserved in LHCII genes of vascular plants, starting at position −415 bp, and the GT1 consensus GRWAAW from −529 bp on, which is found in several light responsive genes of vascular plants (Zhou, 1999). These sequences are however not encoded in the 255 bp fragment of the *LHCBM6* promoter, which was shown to be sufficient to drive light-dependent transcription (Hahn and Kück, 1999). It might therefore be questionable whether these elements are conserved in *C. reinhardtii*. More important, empirical data suggest that under low and medium light, NAB1 expression does not alter much (Berger et al., 2016).

Additionally, PlantRegMap database (http://plantregmap.cbi.pku.edu.cn/; (Jin et al., 2017; Tian et al., 2020)) was used to screen the NAB1 promotor for putative transcription factor binding sites. According to the *in silico* prediction analysis, 12 candidate binding sites of ten transcription factors (TF) were identified for the full (1.5kb) NAB1 promotor upstream sequence before the translation start (Supplementary Table S2 and Supplementary Figures S2 and S3). Among the putative transcription factors, seven different TF families (including SBP, bHLH, C3H, bZIP, Nin-like, CPP and MYB-related superfamilies of transcription factor) were detected, mostly whitin the first 400 bp upstream sequence before the translation start (Supplementary Table S2 and Supplementary Figures S2 and S3). SQUAMOSA promoter binding proteins (SBPs) form a major family of plant/algae-specific transcription factors and play critical roles in regulating flower and fruit development as well as other numerous physiological processes (Kropat et al., 2005; Guo et al., 2008). C3H proteins represent a large family containing zinc finger Cys3His-type motifs, and function most likely as RNA-binding proteins but were also shown to interact with specific DNA sequences under drought stress (Li and Thomas, 1998; Jiang et al., 2014). Nin-like (for nodule inception) family proteins display similarity to transcription factors, and the predicted DNA-binding/dimerization domain identifies and typifies a consensus motif conserved in plant proteins with a function in nitrogen-controlled development (Schauser et al., 1999). CPP-like (cystein-rich polycomb-like protein) proteins are members of a small transcription factor family, which were described to play an important role in development of reproductive tissue and control of cell division in plants (Yang et al., 2008). The MYB-like family of proteins is large, functionally diverse transcription factor family, which had been reported to function in a variety of plant-specific processes and also to interact with other transcription factors (Kirik and Bäumlein, 1996; Ambawat et al., 2013). The basic/helix-loop-helix (bHLH) proteins are a superfamily of transcription factors that are important regulatory components in transcriptional networks in these systems, controlling a diversity of processes from cell proliferation to cell lineage establishment (Toledo-Ortiz et al., 2003). Basic region/leucine zipper motif (bZIP) transcription factor family, which binding site is located far from the translation start (−1007 to - 994 bp, Table S2) had been described to regulate various biological processes and stress responses including pathogen defense, light and stress signaling in plants and algae (Jakoby et al., 2002; Ji et al., 2018).

In summary, this variety of regulating factor, *cis*-regulatory elements as well as responding factors and motifs within the *NAB1* promoter suggests that the *NAB1* gene expression is quite tightly regulated. This conclusion may further imply that in addition to the strong regulation of the activity of the cytosolic repressor NAB1 on protein level (Mussgnug et al., 2005; Wobbe et al., 2009; Blifernez et al., 2011; Berger et al., 2014; Berger et al., 2016), the regulation on gene level also responds to a variety of physiological stressors.

## Footnotes

1 orange: elements involved in copper and hypoxia signalling (curecore; (Quinn et al., 2002; Kropat et al., 2005)); yellow: motifs conferring CO2-responsiveness (Winck et al., 2013); blue: low temperature response elements (ltre; (Jiang et al., 1996; Dunn et al., 1998; Kim et al., 2002)); green: *experimentally determined transcription start site; TATA-box and AT-rich region are putative alternative start sites;

2 grey: 5’UTR in box: putative binding sites for transcription factor (TFbs) (PlantRegMap, http://planttfdb.cbi.pku.edu.cn, (Jin et al., 2017), using default settings)

1 orange: elements involved in copper and hypoxia signalling (curecore; (Quinn et al., 2002; Kropat et al., 2005)); yellow: motifs conferring CO2-responsiveness (Winck et al., 2013); blue: low temperature response elements (ltre; (Jiang et al., 1996; Dunn et al., 1998; Kim et al., 2002)); green: *experimentally determined transcription start site; TATA-box and AT-rich region are putative alternative start sites;

2 grey: 5’UTR in box: putative binding sites for transcription factor (TFbs) (PlantRegMap, http://planttfdb.cbi.pku.edu.cn, (Jin et al., 2017), using default settings)

